# Muscleblind-like proteins use modular domains to localize RNAs by riding kinesins and docking to membranes

**DOI:** 10.1101/2022.07.06.498900

**Authors:** Ryan P. Hildebrandt, Kathryn R. Moss, Aleksandra Janusz-Kaminska, Luke A. Knudson, Lance T. Denes, Tanvi Saxena, Devi Prasad Boggupalli, Zhuangyue Li, Kun Lin, Gary J. Bassell, Eric T. Wang

## Abstract

RNA transport and local translation provide spatial control of gene expression, and RNA binding proteins (RBPs) act as critical adapters in this multi-step process. Muscleblind-like (MBNL) RNA binding proteins, implicated in myotonic dystrophy and cancer, localize RNAs to myoblast membranes and distal neurites through unknown mechanisms. We found that MBNL forms motile and anchored granules in neurons and myoblasts, and selectively associates with kinesins Kif1bα and Kif1c through its zinc finger (ZnF) domains. Other RBPs with similar ZnFs also associate with these kinesins, implicating a motor-RBP specificity code. Live cell imaging and fractionation revealed that membrane anchoring is mediated through the unstructured carboxy-terminal tail of MBNL1. Both kinesin- and membrane-recruitment functions were reconstituted using MBNL-MS2 coat protein fusions. This approach, termed RBP Module Recruitment and Imaging (RBP-MRI), decouples RNA binding, kinesin recruitment, and membrane anchoring functions, while also establishing general strategies for studying multi-functional, modular domains of RBPs.

## Introduction

Cells contain highly organized microenvironments in which mRNA localization and translation are spatiotemporally regulated to influence cell structure and function (Martin and Ephrussi, 2009). This type of regulation is essential in highly polarized and differentiated cell types, such as multinucleated skeletal muscle and neurons (Donlin-Asp et al., 2017). mRNA *cis*-elements, often in 3’ UTRs, recruit *trans*-acting RNA binding proteins (RBPs), facilitating assembly of ribonucleoprotein (RNP) granules competent for transport and local translation (Kiebler and Bassell, 2006). Transport of RNPs is often achieved by association with kinesin motors and travel along microtubules. Indeed, over 40 RBPs are associated with conventional kinesin-1 (KIF5) (Kanai et al., 2004). For example, ZBP1 (IGF2BP1) associates with KIF5 and KIF11 to regulate β-actin mRNA localization and local translation in neurons and fibroblasts, respectively (Hüttelmaier et al., 2005; Sasaki et al., 2010; Song et al., 2015; Wu et al., 2020), and FMRP associates with the kinesin light chain of KIF5 to transport RNA and regulate local translation in an activity-dependent manner (Dictenberg et al., 2008; Ifrim et al., 2015). Recently, RNAs and translational machinery were shown to travel with late endosomes and lysosomes, via adapters such as annexin A11 (Cioni et al., 2019; Liao et al., 2019), implicating membrane-bound vesicular trafficking as yet another localization mechanism. Despite critical roles for all these components to regulate mRNA transport, it remains unclear whether a code exists to specify RBP-motor associations. It is possible that certain kinesin tails may associate with certain RBP domains, facilitating specificity for transport, anchoring, and response to physiological cues. Because many RBPs recognize specific mRNAs via sequence and/or structure, kinesin association and transport behavior may be influenced by RNP composition.

Genetic disruptions to kinesins and RBPs can cause disease. Loss of FMRP causes fragile X syndrome, concomitant with impairment of mRNA localization and local translation (Bassell and Warren, 2008). Mutations in kinesins are linked to Charcot-Marie-Tooth disease (Zhao et al., 2001), hereditary spastic paraplegia (Pennings et al., 2020; Reid et al., 2002), and amyotrophic lateral sclerosis (Nicolas et al., 2018; Wang et al., 2016). In the repeat expansion disease myotonic dystrophy (*dystrophia myotonica*, DM), expanded CTG or CCTG repeats are transcribed into toxic RNA that form intranuclear foci, sequestering and functionally depleting the muscleblind-like (MBNL) family of RBPs (Thornton et al., 2017). MBNLs are deeply conserved throughout metazoa, and mammals encode 3 paralogs: MBNL1, MBNL2, and MBNL3 (Fardaei et al., 2002; Kanadia et al., 2003; Oddo et al., 2016). MBNL1 shows highest expression in heart and muscle, MBNL2 in brain (Charizanis et al., 2012; Kanadia et al., 2003), and MBNL3 in placenta and muscle satellite cells (Poulos et al., 2013). In the nucleus, MBNLs regulate alternative splicing (Jiang et al., 2004; Wang et al., 2012) and polyadenylation (Batra et al., 2014), and loss of MBNL activity in DM causes mis-splicing and several disease symptoms, such as muscle weakness, myotonia, and cardiac arrhythmia (Freyermuth et al., 2016; Fugier et al., 2011; Mankodi et al., 2002). In the cytoplasm, MBNLs also regulate mRNA localization (Adereth et al., 2005). MBNLs enhance association of target RNAs to membranes in myoblasts, particularly those encoding secreted proteins (Wang et al., 2012). In neuronal culture, MBNLs enhance localization of alternative 3’ UTR isoforms to neurites (Taliaferro et al., 2016), and in rat hippocampal neuropil, localized RNAs are enriched for MBNL motifs (Tushev et al., 2018). Indeed, a transgenic mouse model expressing expanded CTG repeats led to reduced levels of cytoplasmic MBNL1 in dendrites prior to missplicing changes (Wang et al., 2017). Additionally, repeat expression in cultured neurons affected neurite outgrowth, which was rescued by co-expression of cytoplasmic, but not nuclear, MBNL1 (Wang et al., 2018). Together, these findings suggest perturbed subcellular distribution of MBNLs and its target RNAs may also drive disease symptoms.

Despite an established role for MBNLs to traffic RNAs in various cell types, we do not understand mechanisms for how MBNL proteins achieve these functions. MBNLs possess two pairs of zinc fingers (ZnFs) separated by an unstructured linker, as well as an unstructured C-terminal domain. Specificity for mRNA targets is achieved via the ZnFs (Lambert et al., 2015; Wagner et al., 2016; Wang et al., 2012), with ZnF1/2 of MBNL1 showing greater specificity for YGCY motifs than ZnF3/4 (Hale et al., 2018). Both MBNL1 and MBNL2 are extensively alternatively spliced, with 9 alternative isoforms encoded by MBNL1 alone. It is unclear whether MBNL proteins use directed transport to move mRNAs, whether they help anchor mRNAs to final destinations, or whether specific isoforms show preferences for these activities.

Here, we have used molecular, cellular, imaging, and computational approaches to elucidate molecular mechanisms for MBNL-mediated RNA localization and membrane anchoring. MBNL1 and MBNL2 form granules exhibiting processive motility in neurons and myoblasts. A kinesin recruitment assay (Bentley et al., 2015; Yang et al., 2016) revealed that Kif1bα and Kif1c tail domains selectively associate with MBNL zinc fingers in an RNA-binding-independent manner. This association results in co-transport of MBNLs and kinesins in live neurons and their co-immunoprecipitation in cell culture. We also found that Tis11d, an RBP with zinc fingers structurally similar to MBNL, also associates with these kinesins, suggesting a structure-function relationship among RBP domains and kinesin tails. Dominant negative expression of specific kinesin tails or functional depletion of MBNL by CTG repeat over-expression, followed by subcellular fractionation and RNA-seq, revealed transcriptome-wide mis-localization of mRNAs in neuronal cell lines and, specifically, depletion of nucleolin mRNA from neurites. Cellular fractionation and live imaging revealed that the unstructured C-terminal tail of MBNL facilitates membrane anchoring and mRNA recruitment to membranes, defining a new function for this domain and providing mechanistic bases for previous observations. We decoupled mRNA binding and kinesin association using a reconstituted synthetic system we term RBP-MRI, highlighting the multi-functional modularity of RBPs, and providing a platform for future studies of other RBP-kinesin pairs. These findings together indicate that certain kinesins can associate with certain RBPs and suggest that a modular kinesin-RBP code may be widespread yet specific, with broad implications for regulation of mRNA trafficking in diverse cell types and tissues.

## Results

### MBNL proteins form motile granules in primary neurons and myoblasts

MBNL1 and MBNL2 mRNA and protein levels rise throughout development, but their protein expression and distribution in cultured primary neurons have not been characterized in detail. To assess the abundance of these proteins in the cytoplasm, where they are presumably involved in localization activities, we examined MBNL localization by immunofluorescence (IF) in mouse cultured primary cortical neurons over time. Both proteins increase in abundance during culture (most dramatically between 8-10 days *in vitro*, DIV), consistent with previous reports of increased mRNA expression (Kanadia et al., 2003). Enrichment of endogenous MBNL2 was observed in the nucleus, whereas MBNL1 showed more cytoplasmic localization (**Fig. S1A**). Similar localization patterns were demonstrated by GFP-MBNL1 fusions, where isoforms lacking exon 5, which contains a nuclear localization signal (NLS), were more cytoplasmic, and those containing exon 5 were more nuclear (**Fig. 1B**, **D**). Higher resolution imaging in 10 DIV neurons revealed MBNL1 and MBNL2 puncta in both axons and dendrites, reminiscent of other RBPs known to regulate mRNA localization (**Fig. 1C**; **Fig. S1C**) (Antar et al., 2004; Zhang et al., 2001).

**Figure 1.**
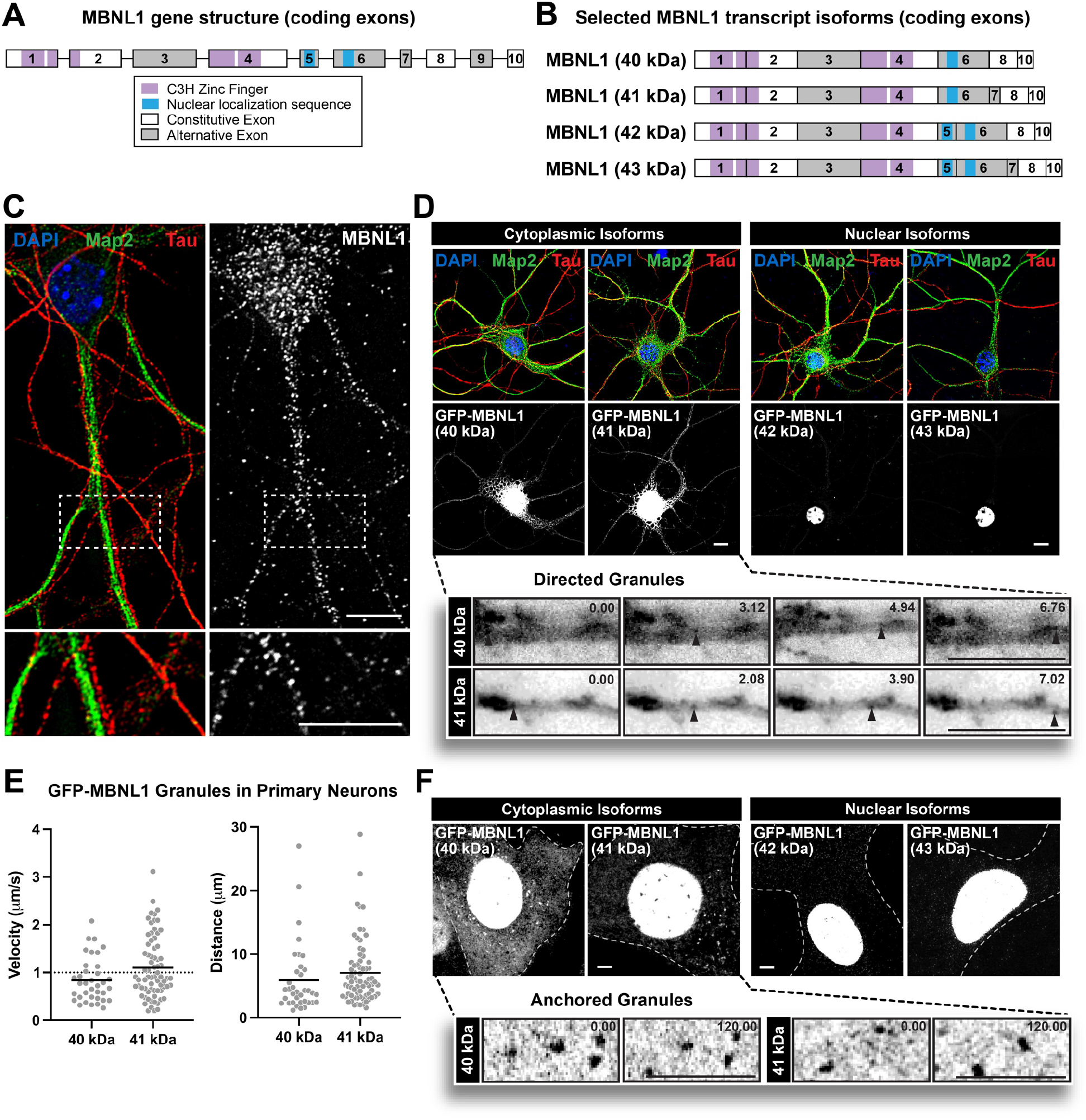
Cytoplasmic MBNL1 granules exhibit directed motion in neurons and anchoring in C2C12 myoblasts. A) Exon and protein domain structure of MBNL1, and B) isoforms studied. C) Representative image showing MBNL1 granules (white) in axons (Tau, red) and dendrites (Map2, green) of 9 DIV cultured primary mouse cortical neurons; nuclei (blue) labeled with DAPI. Scale bar = 10 µm. D) (Above) Representative images of GFP-MBNL1 40, 41, 42, and 43 kDa isoforms (white) in 8 DIV cultured primary mouse cortical neurons; nuclei (blue) labeled with DAPI, axons and dendrites labeled with Tau (red) and Map2 (green), respectively. Scale bar = 10 µm. (Below) Representative time lapse images of motile GFP-MBNL1 40 kDa and 41 kDa granules in live mouse cortical neurons. Time (sec) indicated in top right. Scale bars = 10 µm. F) Quantitation of speeds and distances traveled by cytoplasmic MBNL1 granules (MBNL1-40 kDa – 36 tracks, 1.57 tracks/cell; MBNL1-41 kDa – 71 tracks, 2.37 tracks/cell). Dotted line in velocity chart represents typical speed of microtubule-dependent transport. F) (Above) Representative images of GFP-MBNL1 40, 41, 42, and 43 kDa in C2C12 myoblasts. Dotted lines outline cell boundaries. (Below) Representative time lapse images of anchored GFP-MBNL1 40 kDa and 41 kDa granules in C2C12 myoblasts. Time (sec) indicated in top right. Scale bars = 5 µm.

Granules containing GFP-MBNL1 isoform fusions lacking exon 5 exhibited directed movements in axons and dendrites in 8 DIV neurons (**Fig. 1D**; **Movie S1**). These motile granules moved with velocities consistent with active transport (MBNL1 40 kDa mean velocity ± SEM: 0.84 ± 0.08 μm/sec, MBNL1-41 kDa mean velocity ± SEM: 1.10 ± 0.08 μm/sec; **Fig. 1E**), often displaying long, directed runs (MBNL1-40 kDa mean distance ± SEM: 5.92 ± 0.92 μm, MBNL1-41 kDa mean distance ± SEM: 7.06 ± 0.60 μm, **Fig. 1E**). Again, MBNL2 was predominantly found in the nucleus of primary neurons (**Fig. S1C**), but GFP-MBNL2 isoform fusions lacking exon 5 could be observed in the cytoplasm and showed similar motility (Mean Velocities ± SEM – MBNL2-isoform 1: 1.21 ± 0.09 μm/sec, MBNL2-isoform 3: 1.00 ± 0.08 μm/sec, MBNL2-isoform 4: 1.16 ± 0.05 μm/sec; Mean Distances ± SEM – MBNL2-isoform 1: 7.62 ± 0.75 μm, MBNL2-isoform 3: 6.36 ± 0.73 μm, MBNL2-isoform 4: 8.30 ± 0.57 μm, **Fig. S1D**, **E**).

Live cell imaging of GFP-MBNL1 fusions in C2C12 myoblasts confirmed nuclear versus cytoplasmic localization patterns for isoforms with or without exon 5, respectively. In these cells, most particles were anchored, with rare instances of fast diffusive or directed motion (**Fig. 1F**; **Movie S2**). Together, these observations suggested that MBNL-containing granules are actively transported by molecular motors, yet also possess features that facilitate anchoring to stable, intracellular structures. We subsequently explored mechanisms that might mediate both of these behaviors exhibited by MBNL1, which showed prominent cytoplasmic localization in both muscle and neuronal systems.

### MBNL1 associates with kinesin-3 family members Kif1b⍺ and Kif1c for transport

Given properties suggestive of microtubule-dependent transport, we first sought to identify putative kinesins that might transport MBNL1. We implemented a previously developed split kinesin recruitment assay in which mechanisms of cargo transport can be interrogated by linking various kinesin tails to a kinesin motor or dynein adapter (Bentley et al., 2015; Yang et al., 2016). Fusion of the dynein adapter bicaudal D2 (BicD2) to FKBP and kinesin tails to FRB facilitates dimerization and centrosome recruitment upon the addition of a rapalog molecule (**Fig. 2A**). We confirmed functionality of this system by co-expressing GFP-tagged transferrin receptor (TfR), tdTomato-tagged BicD2-FKBP (BicD2-FKBP), and Myc-tagged Kif1a-FRB or Kif13b-FRB in Neuro2A (N2A) cells. Only the Kif13b tail, and not the Kif1a tail, was able to recruit TfR to the centrosome, consistent with the established ability of Kif13b to carry TfR (**Fig. S2A**) (Bentley et al., 2015). To confirm centrosome targeting, we visualized recruitment of TfR and Kif13b-FRB with BicD2-FKBP, coincident with γ-tubulin staining (**Fig. S2B**). Using this assay, we screened 12 distinct kinesin tails (Kif1a, Kif1bα, Kif1bβ, Kif3a, Kif3b, Kif5a, Kif5b, Kif5c, Kif13a, Kif13b, Kif17, and Kif21b) for their ability to recruit GFP-MBNL1. Kif1bα, but no other kinesin tails (including Kif1bβ, an alternatively spliced isoform from the same gene), strongly recruited a cytoplasmic GFP-MBNL1-40S isoform (**Fig. S2C**) as well as both the 40 kDa and 41 kDa isoforms of MBNL1 (GFP-MBNL1-40 and -41) (**Fig. 2B**, top panels). Recruitment was rapalog-dependent (**Fig. S2D**) and also specific to GFP-MBNL1, but not other proteins such as SMN or TfR (**Fig. S2E-F**). Swapping of the BicD2 adapter for the KIF1A motor domain resulted in rapalog-dependent recruitment of GFP-MBNL1 to plus ends of microtubules at the cell periphery, demonstrating that the association was tail-domain-specific, and not dependent on any specific motor domain (**Fig. 2B**, bottom panels; **Fig. S2G**).

**Figure 2.**
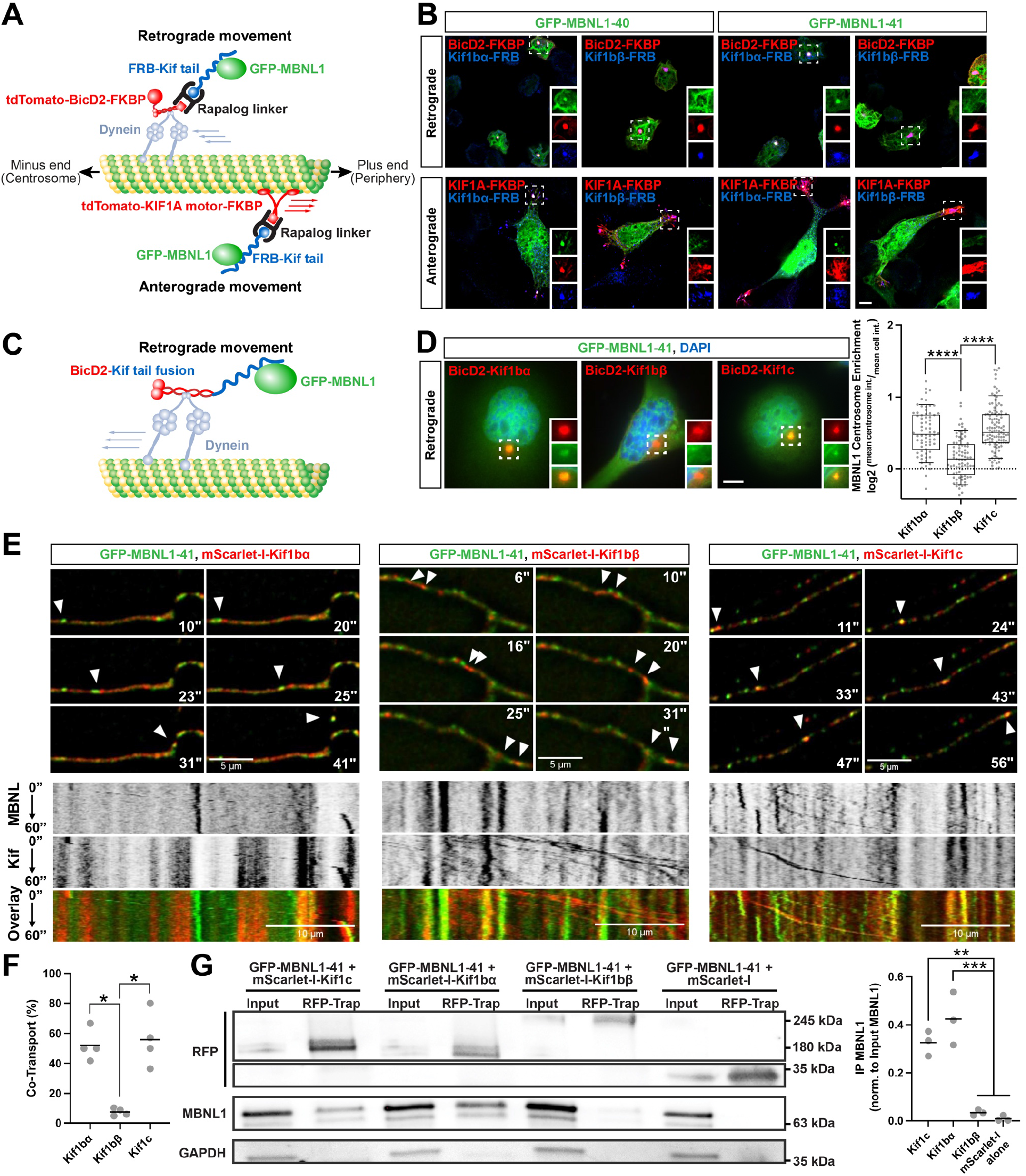
MBNL1 associates, co-transports, and co-immunoprecipitates with Kif1bα and Kif1c. A) Split kinesin recruitment assay used to identify MBNL1-kinesin associations. Cargoes are recruited in a retrograde or anterograde direction in a rapalog-dependent manner, depending on choice of motor complex. B) Representative images showing recruitment of GFP-MBNL1 (green) to the centrosome by Kif1bα, but not Kif1bβ, using Myc-tagged, FRB-kinesin tail fusions (blue) co-expressed with BicD2-FKBP or KIF1A motor-FKBP (red). Scale bar = 10 µm. C) A modified kinesin recruitment assay that uses BicD2-kinesin tail fusions to further characterize MBNL1-kinesin associations. D) Representative images and quantitation of GFP-MBNL1 (green) at the centrosome when co-expressed with BicD2-kinesin tail fusions (red); nuclei (blue) labeled with DAPI (Kif1bα: 70 cells, Kif1bβ: 81 cells, Kif1c: 112 cells; across 3 separate experiments). Scale bar = 10 µm. Bars represent median, box outlines represent upper and lower quartiles, whiskers represent 10th and 90th percentiles. Dotted line represents no enrichment. ****p<0.0001, Two-tailed Mann-Whitney *U* test. E) Representative dual color time lapse images of GFP-MBNL1 (green) and mScarlet-I-kinesins (red) in live 7 DIV primary mouse cortical neurons. Arrows denote co-transported particles of MBNL with Kif1bα (left) and Kif1c (right), whereas Kif1bβ particles lack MBNL (center); time (sec) indicated in top or bottom right. Scale bar = 5 µm. Kymographs are displayed below still frames obtained from videos. Scale bar = 10 µm. F) Fraction of mScarlet-I-kinesin granules showing co-transport with GFP-MBNL1 (across 4 separate experiments). Bars represent mean. *p<0.05, Two-tailed Mann-Whitney *U* test. G) Western blots against RFP, GFP, and MBNL1 following co-immunoprecipitation of mScarlet-Kif fusions with quantitation normalized to total MBNL1-GFP protein (n = 3). Bars denote mean. **p<0.01, ***<0.001, one-way ANOVA with Tukey’s post-hoc test.

We subsequently simplified the centrosome recruitment assay by directly fusing FLAG-tagged BicD2 to Kif1bα and Kif1bβ tail domains to eliminate the need for rapalog addition (**Fig. 2C**). Using this approach, we reconfirmed centrosomal targeting of GFP-MBNL1-41 with Kif1bα but not Kif1bβ (**Fig. 2D**). We additionally tested another kinesin tail that was not included in the initial screen, Kif1c. This kinesin shares ∼58% sequence homology with the Kif1bα tail domain and is highly expressed in the central nervous system and muscle. We developed a quantitative assay to measure the extent of centrosome recruitment (**Fig. S3**), and showed that Kif1bα and Kif1c (log2 mean centrosome enrichment: 0.506 vs. 0.560, **Fig. 2D**) recruit GFP-MBNL1 to the centrosome more strongly than Kif1bβ (log2 mean centrosome enrichment: 0.149, ****p<0.0001, two-sided Mann-Whitney *U* test, **Fig. 2D**).

The centrosome recruitment assay decouples kinesin tails from their motor domains and is not designed to assess co-transport of Kifs and MBNL1. To investigate this further, we turned to primary neurons. To confirm physiological co-incidence of MBNL1 and full-length versions of the candidate Kifs, we performed IF on 12 DIV neurons transduced with AAV expressing HA-MBNL1-41 (**Fig. S4A**). 3D quantitative colocalization analysis of select neurites supported previous findings; endogenous Kif1bα and Kif1c both displayed significantly higher levels of colocalization with HA-MBNL1 than endogenous Kif1bβ with HA-MBNL1 (**Fig. S4B**). While this confirmed static colocalization of MBNL with specific Kifs, we subsequently investigated co-transport through live cell imaging of GFP-MBNL1-41 and mScarlet-tagged Kif1bα, Kif1c or Kif1bβ in DIV 7 primary cortical neurons. Approximately 50-60% of mScarlet-Kif1bα and mScarlet-Kif1c proteins showed long, processive runs (>2 µm) together with GFP-MBNL1 (*p<0.05, two-sided Mann-Whitney *U* test, **Fig. 2E, F**), but <10% of mScarlet-Kif1bβ proteins showed co-transport (p>0.05, two-sided Mann-Whitney *U* test, **Fig. 2E, F**). To confirm these observations biochemically and to test the strength of these associations, we co-expressed mScarlet-Kif proteins with GFP-MBNL1-41 in N2A cells and performed co-immunoprecipitations using an RFP-trap antibody. GFP-MBNL1 showed ∼10-fold enrichment in the presence of Kif1bα and Kif1c as compared to Kif1bβ, which showed less than 2-fold enrichment relative to mScarlet alone, as assessed by western blot (***p<0.001, **p<0.01, p>0.05, one-way ANOVA with Tukey’s test, **Fig. 2G**). In summary, these experiments revealed a selective and functional association between MBNL1 and specific kinesin tails that exists as motile particles in live neurons and persists through biochemical manipulation.

### Expression of repeats and dominant negative kinesin tails causes mis-localization of MBNL-targeted mRNAs

We hypothesized that localization of mRNAs might be perturbed upon either MBNL1 depletion or kinesin motor perturbation. We used a neurite fractionation assay to measure mRNA localization by plating N2A cells onto semi-permeable (1 µm sized pores) membranes to facilitate neurite growth and separation, and at the same time transiently introduced plasmids expressing 480 CTG repeats or dominant negative kinesin tails with no motor domains. Neurite and cell body fractions were validated using antibody markers for each compartment (see Methods). These perturbations either functionally deplete MBNL activity or kinesin dependent transport, respectively (**Fig. 3A**). Cell body and neurite RNA was then subjected to RNA-seq (TPMs for all transcripts in **Table S1**). Depletion of MBNL was induced by expression of 480 CTG repeats to measure mis-localization of MBNL target transcripts. mRNAs containing a greater number of MBNL1 CLIP tags/kilobase in their UTRs, as previously determined (Wang et al., 2012), showed stronger shifts away from neurites relative to soma, as compared to control cells transfected with a DMPK fragment encoding 0 CTG repeats (**Fig. 3B**; **Table S2**). These results mimic previous observations of shifts in MBNL target mRNAs away from neurites following MBNL knockdown (Taliaferro et al., 2016), but use CTG repeats to deplete MBNL activity rather than siRNA. Upon comparison to the kinesin dominant negative conditions, a modest ∼2.6-fold enrichment in overlap was observed between those mRNAs mis-localized away from neurites upon CTG480 expression (relative to CTG0) versus mRNAs similarly mis-localized upon Kif1bα tail over-expression (relative to Kif1bβ over-expression). A similar ∼2.4-fold enrichment in overlap was observed when comparing mRNAs mis-localized upon CTG480 expression to those mis-localized upon Kif1c tail over-expression (**Fig. 3C**).

**Figure 3.**
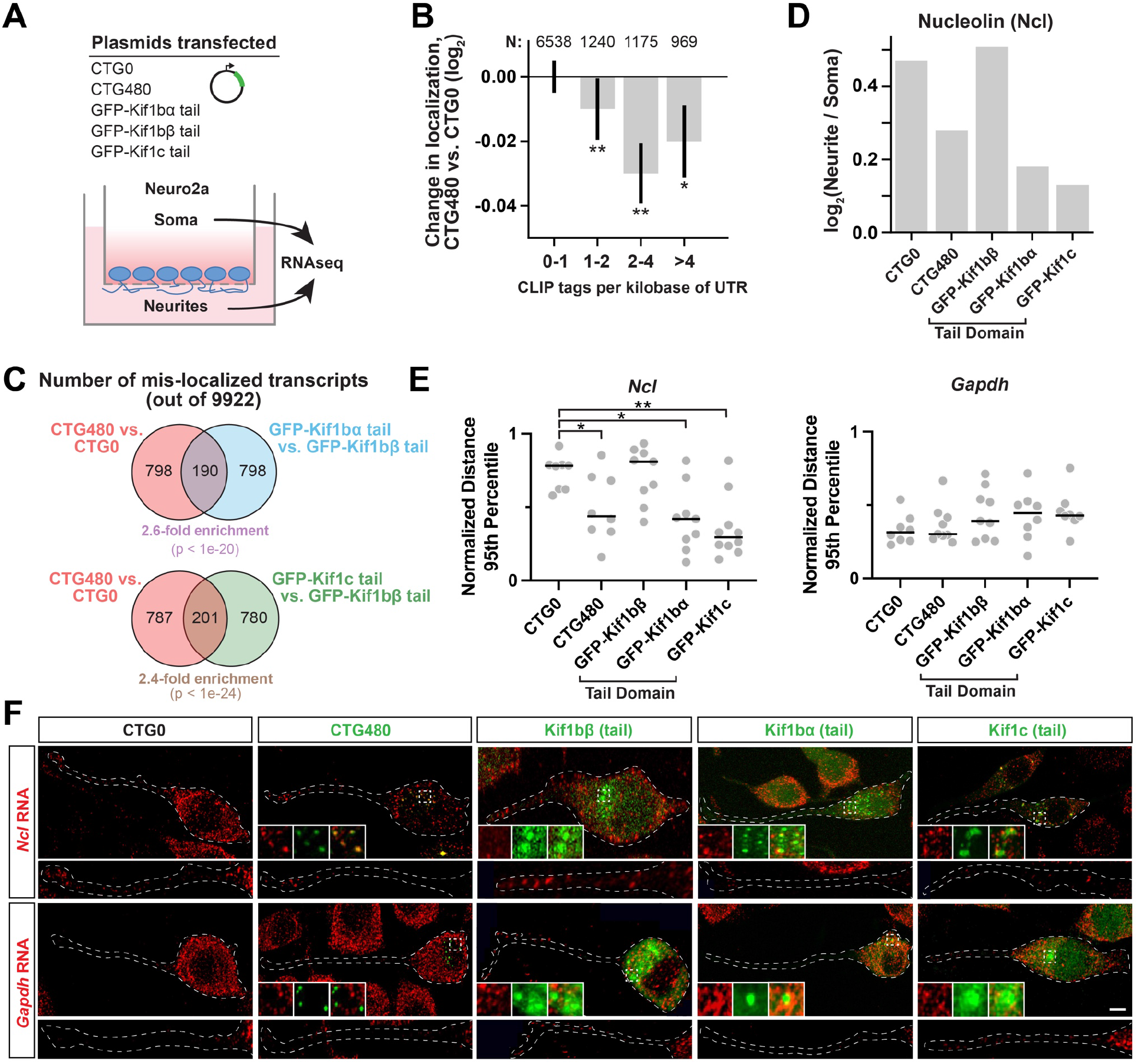
Depletion of MBNL or expression of kinesin tails leads to mis-localization of RNA targets. A) Experimental design used to deplete MBNLs or over-express kinesin tail dominant negatives in differentiated N2A cells, followed by fractionation of soma and neurites. B) Bars showing change in localization of mRNAs from neurite to soma in N2A expressing CTG480 as compared to CTG0, as a function of CLIP tag density per kilobase of 5’ and 3’ UTR. *p<0.01, **p<0.005, rank-sum test. C) Venn diagram showing overlap of mRNA targets mis-localized from neurite to soma upon perturbation by CTG480 relative to CTG0 or Kif1bα/Kif1c tails relative to Kif1bβ tail. P values computed by Fisher’s Exact test, and enrichment of overlap was computed assuming independence between conditions. D) The ratio of nucleolin mRNA between neurite and soma across conditions, as assessed by RNAseq. E) Distances of *Ncl* and *Gapdh* HCR FISH spots from the soma of each cell across conditions. Distances were normalized as a fraction of total neurite length, and the 95th percentile of distances within each cell is plotted as an individual gray circle. Each cell (Ncl – CTG0: 8 cells, CTG480: 8 cells, Kif1bβ: 9 cells, Kif1bα: 9 cells, Kif1c: 10 cells; Gapdh – CTG0: 8 cells, CTG480: 9 cells, Kif1bβ: 9 cells, Kif1bα: 8 cells, Kif1c: 8 cells) contained >150 *Ncl* spots and >300 *Gapdh* spots. Bars represent the median across cells. *p<0.05, **p<0.01, Two-tailed Mann-Whitney U test. F) Representative HCR FISH images for Ncl and Gapdh (red) with CTG repeats or dominant negative kinesin tails (green). Scale bar = 10 µm.

The mRNA encoding nucleolin (Ncl) was mis-localized upon CTG480, Kif1bα tail, and Kif1c tail over-expression (**Fig. 3D**). Nucleolin, an RNA binding protein, inhibits axonal growth of sensory neurons through localization of importin β1 RNA and has shown to be present in axons, potentially playing a role to sense and regulate cell size (Perry et al., 2016). MBNL1 CLIP tags in the vicinity of YGCY motifs are present in the UTRs and coding sequence of the *Ncl* transcript (**Fig. S5A**), which prompted us to confirm subcellular localization by HCR RNA FISH. Indeed, *Ncl* mRNA spots were depleted from neurites upon CTG480 over-expression, and *Ncl* mRNAs co-localized with dominant negative Kif1bα and Kif1c tail proteins in the cytoplasm (**Fig. 3F**). Upon quantitation, the 95th percentile of *Ncl* HCR FISH spot distances, normalized as a percentage of total neurite length from the cell body centroid, was decreased upon CTG480, Kif1bα, or Kif1c tail over-expression, but not upon CTG0 or Kif1bβ overexpression, while *Gapdh* localization was unaffected in all conditions (*p<0.05, **p<0.01, two-sided Mann-Whitney *U* test, **Fig. 3E**). *Ncl* mRNA mis-localization was assessed in another neuronal cell line, CAD, in response to similar perturbations. Similar co-localization of *Ncl* mRNA with Kif1bα and Kif1c was observed, along with striking co-localization with intranuclear repeat foci and significantly decreased spot distance (**Fig. S5B, C**). Together, these data suggest that MBNL1 and specific kinesins may play roles to localize a number of mRNAs to neurites, and that localization of Ncl mRNA to neurites requires MBNL1 and Kif1bα or Kif1c, with implications for neuronal development and function.

### Association of MBNL1 with kinesin motors requires intact zinc fingers

We hypothesized that specific domains of MBNL1 might be responsible for kinesin association. First, RNA dependency was tested by using previously established <u>R</u>NA Interaction <u>M</u>utants (RIMs) that disrupt interactions of MBNL1 zinc fingers (ZnFs) with GC dinucleotides, abolishing RNA binding yet preserving ZnF structure (Purcell et al., 2012). We mutated all 4 ZnFs to abolish RNA binding. Co-expression of GFP-tagged MBNL1-41-RIM with FLAG-tagged BicD2-Kif1c yielded a similar level of centrosome enrichment as compared to wild-type GFP-MBNL1-41, suggesting that kinesin association does not require RNA binding activity (log2 mean centrosome enrichment: 0.556 vs. 0.497, p>0.05 by two-sided Mann-Whitney *U* test, **Fig. 4B**, **C**). The non-ZnF domains of MBNL1 lack defined structure as predicted by AlphaFold (**Fig. 5A**, top) (Jumper et al., 2021; Mirdita et al., 2022), and roles of unstructured domains in other RBPs have been implicated in protein-protein interactions, liquid-liquid phase separation, and stress granule formation, among other functions (Van Treeck and Parker, 2018). The exon 3 linker between ZnF pairs confers flexibility for enhanced RNA binding and splicing regulation (Tran et al., 2011). While the C-terminal tail was initially proposed to contain a trans-membrane domain (Miller et al., 2000), it is now understood to contain a bipartite NLS sequence within exon 6 and a dimerization domain within exon 7 (Tabaglio et al., 2018; Tran et al., 2011). Presence of this C-terminal tail domain enhances formation of stable CUG repeat foci (Arandel et al., 2022). To test the ability of unstructured domains to mediate kinesin association, we generated mutants lacking both exon 3 and the C-terminal tail (MBNL1-Δ3, ΔC RIM) and co-expressed them with BicD2-Kif1c. This protein exhibited a more cytoplasmic distribution compared to the 41 kDa isoforms and, notably, an even stronger enrichment at centrosomes, potentially explained by its increased abundance in the cytoplasm (log2 mean centrosome enrichment: 0.724, ****p<0.0001 by two-sided Mann-Whitney *U* test, **Fig. 4B**, **C**). Therefore, the unstructured domains appear dispensable for kinesin dependent transport.

**Figure 4.**
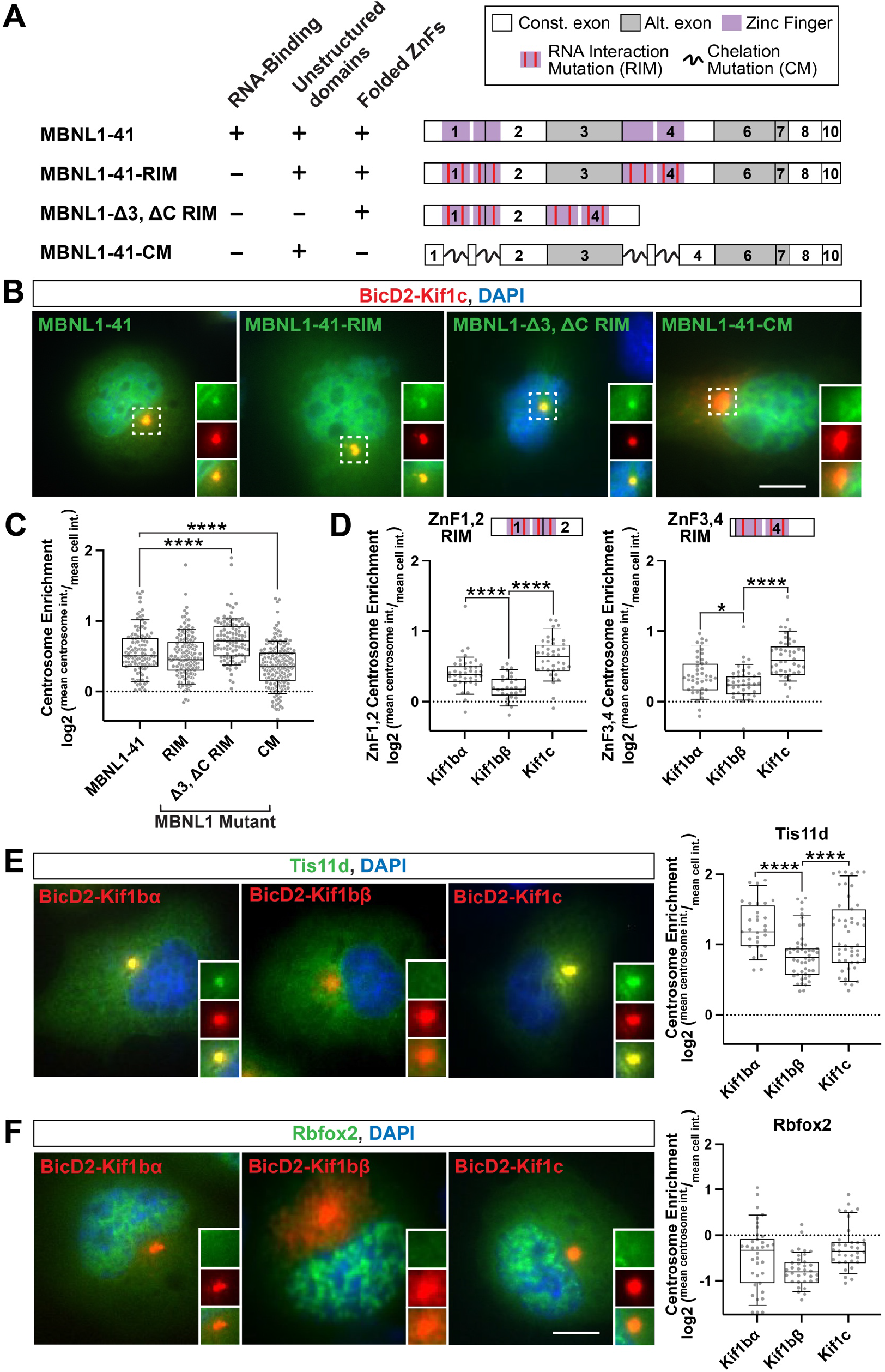
Zinc fingers of MBNL1 are necessary and sufficient to associate with kinesins in an RNA binding-independent manner, and structurally similar zinc fingers in other RBPs associate with the same kinesins. A) Schematic of GFP-MBNL1 mutants tested. B) Representative images of centrosome enrichment for each mutant (green) as tested with BicD2-Kif1c tail fusions (red). Nuclei labeled with DAPI (blue). C) Quantitation of centrosome enrichment for each mutant. MBNL1-41: 112 cells, MBNL1-41-RIM: 146 cells, MBNL1-Δ3, ΔC RIM: 106 cells, MBNL1-41-CM: 182 cells; across 4 separate experiments. D) Quantitation of centrosome enrichment for GFP-ZnF1/2 RIM and GFP-ZnF3/4 RIM mutants using BicD2 kinesin tail fusions. MBNL1-ZnF1,2 RIM – Kif1bα: 39 cells, Kif1bβ: 29 cells, Kif1c: 40 cells; MBNL1-ZnF3,4 RIM – Kif1bα: 42 cells, Kif1bβ: 39 cells, Kif1c: 44 cells; across 3 separate experiments. E) Representative images and quantitation of centrosome enrichment for Tis11D (green) with BicD2 kinesin tail fusions (red). Kif1bα: 29 cells, Kif1bβ: 49 cells, Kif1c: 49 cells; across 3 separate experiments. F) Representative images and quantitation of centrosome enrichment for Rbfox2 (green) with BicD2 kinesin tail fusions (red). Kif1bα: 36 cells, Kif1bβ: 35 cells, Kif1c: 36 cells; across 3 separate experiments. Bars represent median, box outline represent upper and lower quartiles, whiskers represent 10th and 90th percentiles. Dotted line represents no enrichment (=1). Scale bars = 5 µm. ****p<0.0001, **p<0.01, Two-tailed Mann-Whitney *U* test. Nuclei labeled with DAPI (blue).

**Figure 5.**
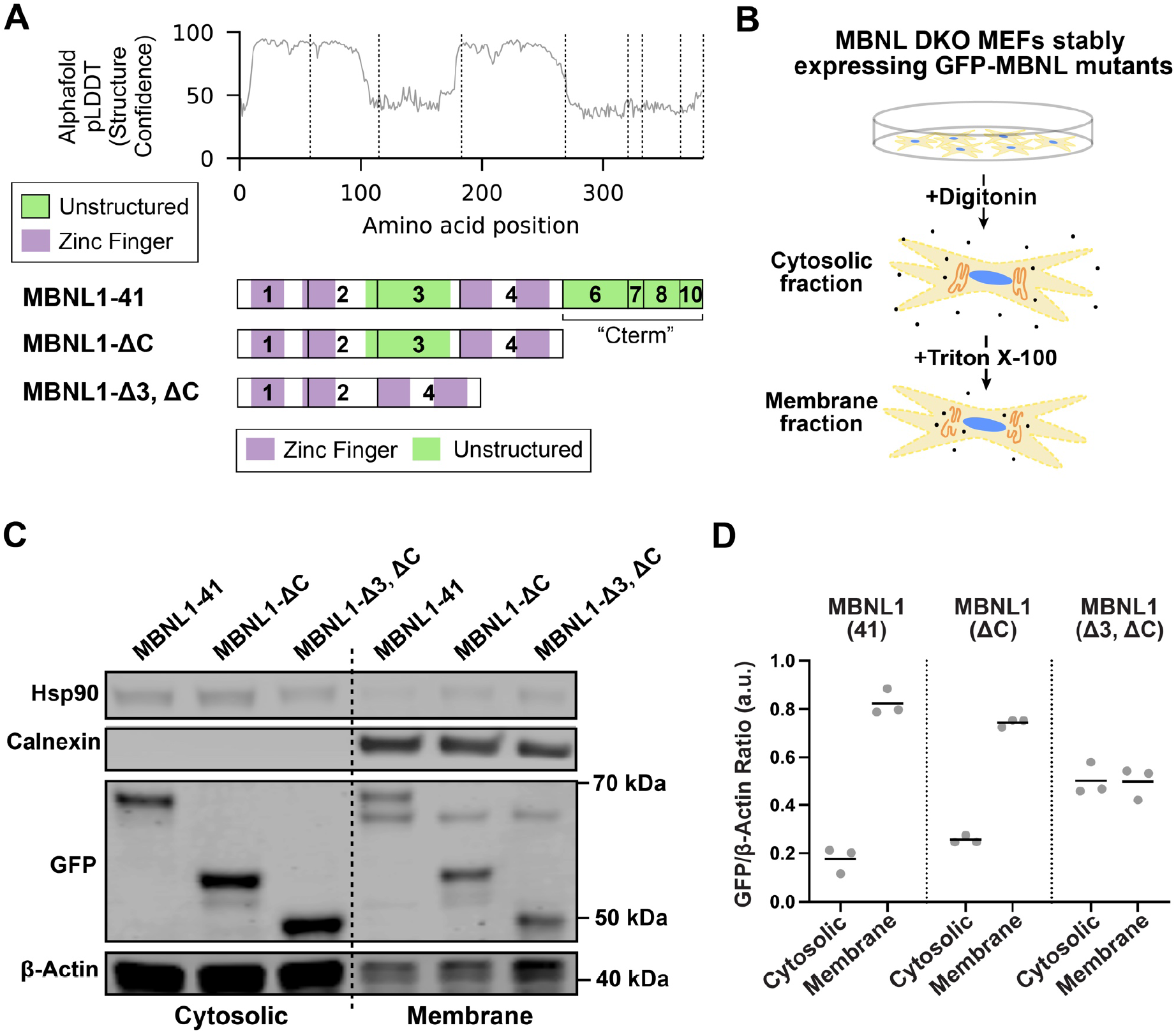
Unstructured domains of MBNL1 confer membrane association properties. A) Structure confidence as assessed by Alphafold for the 41 kDa isoform of MBNL1; this metric is aligned to the exon structure below, and additional MBNL1 mutants which lack exon 3 and/or the C-terminal tail are also illustrated. B) Schematic of detergent-based subcellular fractionation approach in MEFs. C) Representative Western blot of GFP-MBNL1 mutants following subcellular fractionation of cytosolic and membrane-associated compartments. Hsp90 shows enrichment in the cytosolic fraction; Calnexin the membrane fraction. D) Quantitation of GFP-MBNL1 concentration in each compartment as normalized to β-actin loading control (n = 3).

Because the unstructured domains did not appear to mediate kinesin association, we then tested whether the ZnFs themselves possess these functions. We used previously generated chelation mutants (CMs) that contain point mutations to cysteine and histidine residues, preventing chelation of zinc ions and thus proper ZnF folding (Purcell et al., 2012). GFP-tagged MBNL1-41-CM showed significantly decreased recruitment at centrosomes when co-expressed with BicD2-Kif1c, revealing that indeed the ZnF domains are important for MBNL-kinesin interactions (log2 mean centrosome enrichment: 0.356, ****p<0.0001 by two-sided Mann-Whitney *U* test, **Fig. 4B**, **C**). Isolated ZnF pairs (ZnF1,2 and ZnF3,4) alone showed significant recruitment with BicD2-Kif1bα and BicD2-Kif1c compared to BicD2-Kif1bβ, although recruitment with Kif1c was slightly stronger (**Fig. 4D**).

These observations prompted questions regarding whether ZnFs with structure similar to those of MBNL1 might also associate with the same kinesin tails. Another RBP with structurally similar ZnFs is Tis11d (Teplova and Patel, 2008); therefore, we tested Tis11d in our centrosome recruitment assay. Strikingly, we observed similar patterns of kinesin tail specificity, with Kif1c and Kif1bα recruiting efficiently, but not Kif1bβ (log2 mean centrosome recruitment Kif1bα vs. Kif1bβ vs. Kif1c – 1.235 vs. 0.826 vs. 1.131, ***p<0.001, **<0.01, two-sided Mann-Whitney *U* test, **Fig. 4E**). This same pattern of recruitment was not observed with Rbfox2, which contains 3 RNA recognition motifs (RRMs) rather than ZnFs; this RBP exhibited little association with any of the 3 kinesin tails tested (**Fig. 4F**). Together, these findings implicate a novel role for MBNL1 ZnFs to associate with specific kinesins for directed transport in an RNA binding-independent manner and suggest a structural code that may underlie how trafficking of mRNAs is controlled by specific RBP-kinesin associations.

### Unstructured domains of MBNL1 enhance association with membranes

We next addressed potential roles of the unstructured domains of MBNL in mRNA localization. Previously, MBNLs were established to localize mRNAs to endoplasmic reticulum membranes for local translation of secreted proteins (Wang et al., 2012). Here, observations of immobile granules in C2C12 myoblasts suggest a membrane anchoring function (**Fig. 1F**; **Movie S2**). We directly tested whether unstructured domains of MBNL1 might facilitate membrane docking and association by studying GFP-tagged, truncated versions of MBNL1 lacking either the C-terminal tail, or the C-terminal tail and exon 3 (ΔC or Δ3, ΔC mutants, **Fig. 5A**). These truncated MBNL1 proteins, along with full-length MBNL1, were stably introduced into mouse embryonic fibroblasts (MEFs) lacking endogenous Mbnl1 and Mbnl2, to preclude any potential piggybacking on target mRNAs that might localize to membranes via endogenous MBNL proteins. These cells were fractionated using digitonin and Triton X-100 to yield cytosolic and membrane-associated compartments, respectively (**Fig. 5B**). Successful separation was confirmed by enrichment of Hsp90 and calnexin in cytosolic and membrane fractions, respectively (**Fig. 5C**). All GFP-tagged MBNL1 proteins were detected in both fractions, and quantitation of GFP-MBNL1 relative to β-actin protein showed greatest membrane enrichment for full-length MBNL1, decreased enrichment upon removal of the C-terminal tail, and the least enrichment upon removal of both C-terminal tail and exon 3 (**Fig. 5D**). These observations suggest that the unstructured domains of MBNL1 contribute to membrane docking, with implications for mRNA localization and local translation.

### Recruitment of a synthetic mRNA to membranes by heterologous reconstitution of MBNL unstructured domains

While experiments above support association of MBNL unstructured domains with membranes, they do not prove a direct role in RNA targeting. To test this possibility, we developed C2C12 myoblasts stably expressing a ponasterone-inducible 45xMS2 RNA reporter, and integrated various tetracycline-inducible fusions of MBNL1 domains to MS2 coat protein (MCP) and HaloTag (MCP-Halo) for live cell imaging of RNP granules (**Fig. 6A**, bottom) (Bertrand et al., 1998). C2C12 myoblasts expressing MCP-Halo fusions without MS2 RNA reporter exhibit diffuse signal, whereas co-expression of the MS2 reporter yields visible motile granules in the cytoplasm (**Movie S5**). The utility of this approach, which we term <u>RBP</u> <u>M</u>odule <u>R</u>ecruitment and Imaging (RBP-MRI), lies in the fact that each visible particle contains multiple copies of the RBP of interest, thus providing a readout biased towards behavior of that RBP. We obtained time lapse images of C2C12 myoblasts at 10 frames per second for 1 minute and analyzed particle tracks using the Fiji plugin TrackMate (representative tracks shown in **Fig. 6B**; **Movie S2**) (Tinevez et al., 2017).

**Figure 6.**
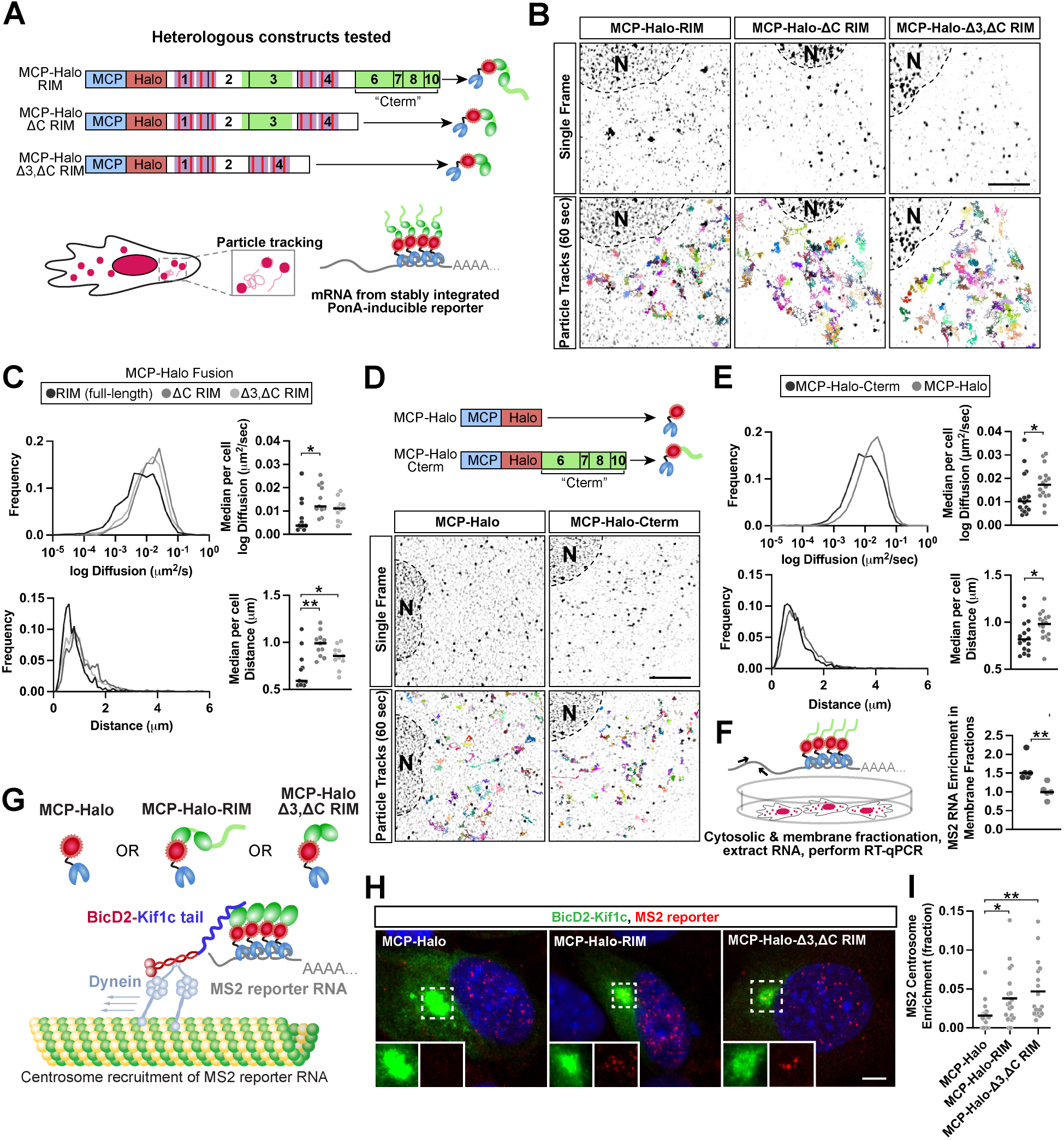
Heterologous reconstitution of domains from MBNL to localize RNAs in a kinesin- or membrane anchoring-dependent manner. A) MS2 reporter system used to image single RNA molecules and reconstitute activities of MBNL domains to localize RNAs by fusion to MCP-Halo. Fusions generated include full-length MBNL1 or truncated MBNL1 without C-terminal tail and/or exon 3; ZnFs in these contexts were all RIM mutants incapable of binding RNA. B) Particle tracks (multi-colored) from videos of C2C12 myoblasts stably expressing MCP-Halo-RIM, MCP-Halo-ΔC RIM, or MCP-Halo-Δ3, ΔC RIM. C) Inferred ensemble log diffusion coefficients (µm^2^/s) and maximum distance (µm) traveled for particles in each condition; MCP-Halo-RIM: 1233 particle tracks, MCP-Halo-ΔC RIM: 1039 particle tracks, MCP-Halo-Δ3, ΔC RIM: 934 particle tracks. Median per cell log diffusion coefficients (µm^2^/s) and maximum distance (µm); 11 cells per condition. D) Schematic and particle tracks (multi-colored) from videos of C2C12 myoblasts stably expressing MCP-Halo or MCP-Halo-Cterm. E) Inferred ensemble log diffusion coefficients (µm^2^/s) and maximum distance (µm) traveled for particles in each condition; MCP-Halo: 1205 particle tracks, MCP-Halo-Cterm: 885 particle tracks. Median per cell log diffusion coefficients (µm^2^/s) and maximum distance (µm); 16 cells per condition. F) Enrichmentof MS2 reporter mRNA as assessed by RT-qPCR from membrane and cytosolic fractions in the presence of MCP-Halo or MCP-Halo-Cterm. G) Schematic of MS2 reporter system used to reconstitute kinesin-dependent transport of RNA via MBNL1 zinc fingers. H) Representative images of HCR FISH (red) against the MS2 mRNA reporter in the presence of BicD2-Kif1c (green) and Halo-MCP fusions. Nuclei labeled with DAPI (blue). I) Quantitation of the fraction of SunTag-MS2 RNA molecules present at the centrosome relative to the whole whole cell across each condition. Scale bars = 5 µm. *p<0.05, **p<0.001, Two-tailed Mann-Whitney *U* test.

To assess contributions of the unstructured C-terminal tail and/or exon 3 to RNP dynamics, MCP-Halo-MBNL1-RIM was first compared to MCP-Halo-MBNL1-ΔC RIM and MCP-Halo-MBNL-Δ3, ΔC RIM (**Fig. 6A**, top). The use of RIM mutants and the MS2-MCP system decouples observations of MBNL1-mediated RNA localization dynamics from the RNA binding activity of MBNL1 ZnFs. Analyses of ensemble particle diffusion coefficients and maximum distance traveled showed that particles containing the MBNL1 C-terminal tail travel shorter distances and are least diffusive (median ensemble diffusion: 6.7×10^-3^ μm^2^/s, median ensemble distance: 0.689 μm, **Fig. 6C**). The presence of exon 3 did not appreciably modulate particle behavior beyond the contribution of the C-terminus (median ensemble diffusion of ΔC RIM: 1.47×10^-2^ μm^2^/s, Δ3, ΔC RIM: 1.16×10^-2^ μm^2^/s; median ensemble distance of ΔC RIM: 0.988 μm, Δ3, ΔC RIM: 0.879 μm, **Fig. 6C**). We next tested whether the C-terminal tail alone, decoupled from any ZnFs, might be sufficient to enhance anchoring dynamics (**Fig. 6D**). MCP-Halo alone showed diffusive, Brownian motion, but MCP-Halo fused to the C-terminal tail showed much more confined dynamics (median ensemble diffusion of +C-term: 1.10×10^-2^ μm^2^/s, MCP-Halo: 2.31×10^-2^ μm^2^/s; median ensemble distance of +C-term: 1.15 μm, MCP-Halo: 1.55 μm, **Fig. 6E**). Taken together, these results show that the MBNL1 C-terminal domain is necessary and sufficient for restricted mobility and RNA anchoring.

These live imaging experiments illustrate how the C-terminal domain restricts RNP motion, but they do not directly link these changes in mobility to membrane association of target RNAs. To confirm this hypothesis, we performed subcellular fractionation, similar to experiments in Fig. 5, and assessed the role of the MBNL1 C-terminal domain to influence MS2 RNA abundance in membrane and cytosolic compartments using RT-qPCR (**Fig. 6F**). The presence of the C-terminal domain promoted ∼60% greater association of MS2 reporter RNA with the Triton-sensitive membrane fraction relative to the cytosol, as compared to a Gapdh control mRNA (**p<0.01, two-tailed Mann-Whitney *U* test, **Fig. 6F**). These observations suggest that the C-terminal tail anchors MBNL1 and its target mRNAs to membranes, consistent with slower RNP dynamics as visualized in live cells. Analogous to the role of zinc fingers to mediate kinesin motor association, the C-terminal domain performs these functions in an RNA binding-independent manner.

### Kinesin tail-dependent transport of a synthetic mRNA by heterologous reconstitution of MBNL ZnFs

In previous experiments, we showed that MBNL1 ZnFs associate with specific kinesin tails, as supported by co-transport in cells and by co-immunoprecipitation in a detergent-resistant manner. However, this does not prove that these associations facilitate RNA transport. To demonstrate this, we again turned to our RBP-MRI approach. To assess whether MBNL ZnFs can localize RNAs in a kinesin-dependent manner, we co-expressed a BicD2-Kif1c tail fusion, which pulls cargoes to the centrosome, with stably integrated tet-inducible MCP-Halo, MCP-Halo-MBNL1 (RIM), or MCP-Halo-MBNL1 (Δ3, ΔC RIM) fusions (**Fig. 6G**). We performed HCR FISH against the MS2 reporter RNA and counted RNA molecules at the centrosome and in the entire cell (**Fig. 6H**). Two to three-fold greater MS2 RNA molecules localized to the centrosome in the presence of both MCP-Halo-RIM fusions, as compared to MCP-Halo alone (**Fig. 6I**). These observations demonstrate that MBNL ZnFs are sufficient to localize RNAs to distinct subcellular locations in a kinesin-dependent manner, independent of ZnF RNA binding activity. Overall, this heterologous system has allowed us to decouple and characterize distinct activities of modular MBNL protein domains, and synthetically reconstitute them in a variety of systems.

### MBNLs regulate the biogenesis of their kinesin transport partners

Importantly, Kif1bα and Kif1bβ are alternative isoforms generated from the *Kif1b* gene; Kif1bα is generated by use of an alternative last exon and earlier polyadenylation site relative to Kif1bβ (**Fig. 7A**). MBNL is well established to regulate alternative splicing, alternative last exon usage, and alternative polyadenylation in the nucleus. To our surprise, examination of previously generated datasets (Oddo et al. 2016, Wang et al., 2019) show that the ratio of Kif1bα to total Kif1b changes upon MBNL depletion. Overall Kif1bα/KIF1Bα levels relative to total Kif1b/KIF1B are decreased in mouse embryonic fibroblasts lacking Mbnl1 and Mbnl2 (DKO), and in human DM1 tibialis anterior biopsies (**Fig. 7B**, **C**). These findings suggest that MBNL regulates the production of specific kinesin isoforms with which it subsequently associates in the cytoplasm. Thus, we find that not only does MBNL mediate several distinct steps in RNA localization, including RNA binding, kinesin engagement, and membrane anchoring, but that it also regulates the abundance of the specific motors with which it interacts (**Fig. 7D**). In principle, this molecular circuitry may allow the cell to appropriately match supply and demand of a specific motor by using the same RBP in both the nucleus and cytoplasm.

**Figure 7.**
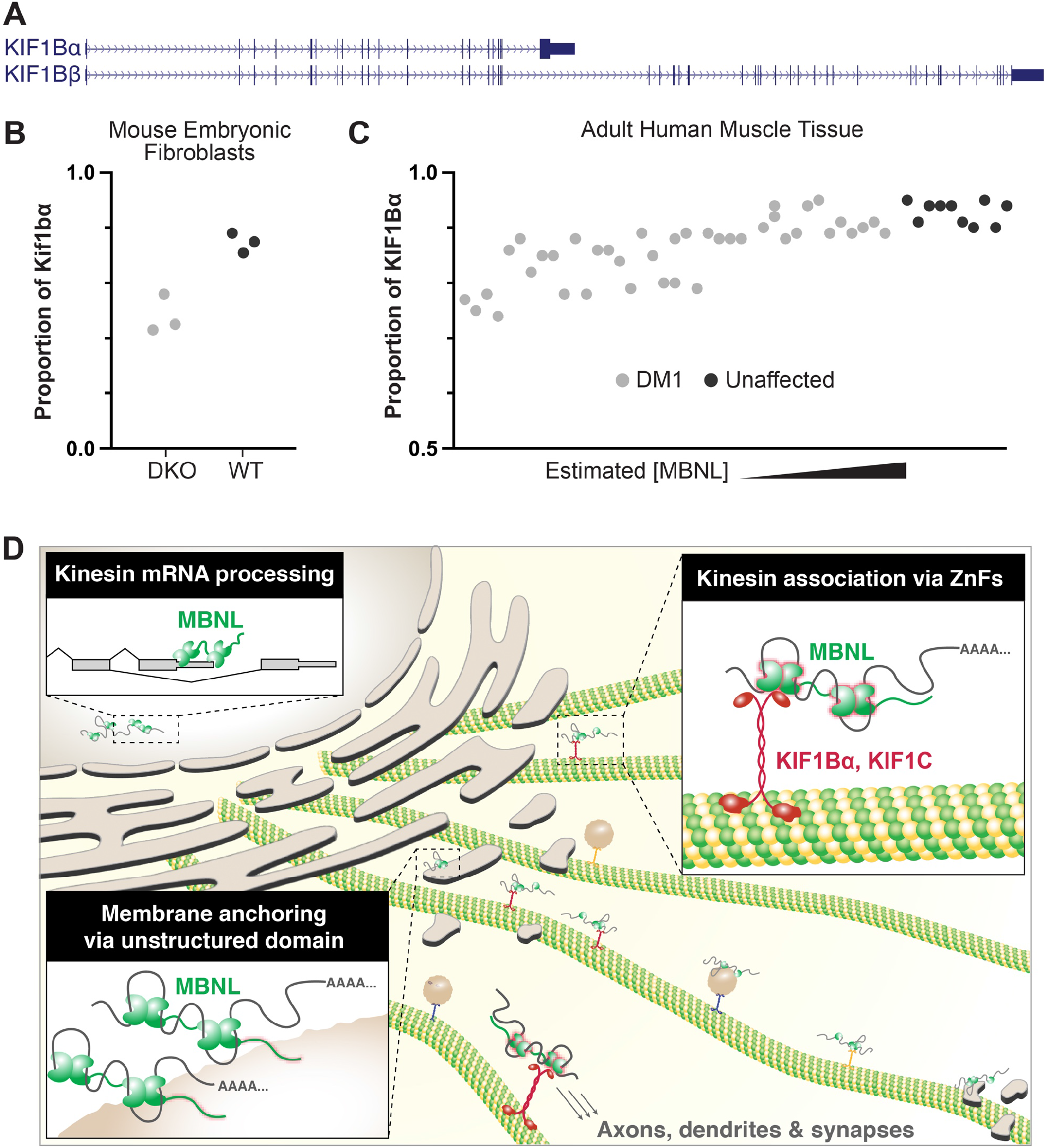
MBNL1 regulates the production of kinesin tails with which it associates in the cytoplasm. A) Representation of alternative last exon usage in the KIF1B gene, which yields two isoforms. B) Proportion of Kif1bα vs. total Kif1b in Mbnl1/2 DKO MEFs and C) KIF1Bα vs total KIF1B in human DM1 tibialis (right). D) Model for how MBNLs regulate production of kinesin tails with which they associate in the cytoplasm to travel along microtubules and anchor at membrane destinations.

## Discussion

Highly differentiated cells use molecular motors and RBPs to localize mRNAs to subcellular compartments to generate and maintain local protein gradients, but the specific interactions between these molecules necessary for mRNA localization are not clear. This is particularly apparent in neurons (Hafner et al., 2019), muscle (Denes et al., 2021; Pinheiro et al., 2021), heart (Scarborough et al., 2021), intestinal epithelium (Moor et al., 2017), and other cell types. There are thousands of localized mRNAs, over 1000 RNA binding proteins, and over 40 kinesin motor proteins in the human genome, producing a large number of potential combinations. Immunoprecipitation-based methods have identified putative binding partners for a number of kinesins, most notably, conventional kinesin-1, kinesin-2, kinesin-3 family members, and others such as KIF21, KIF26A, and KIF13B (Asaba et al., 2003; Horiguchi et al., 2006; Setou et al., 2002; Takeda et al., 2000; Weng et al., 2018; Zhou et al., 2009). Some RBPs appear on these lists, but a surprisingly small number of kinesin-RBP pairs with functional interactions have been established thus far. FMRP, ZBP1, hnRNP Q, Pur-α, SFPQ, Staufen1, egalitarian, and tropomyosin are RBPs that have been associated with specific kinesins in mammalian and *Drosophila* systems, but the sheer number of motors and cargoes implies that we have only explored a small subset of all possible interactions (Bannai et al., 2004; Brendza et al., 2000; Davidovic et al., 2007; Dictenberg et al., 2008; Fukuda et al., 2020; Gáspár et al., 2017; Kanai et al., 2004; McClintock et al., 2018; Ohashi et al., 2002; Song et al., 2015; Wu et al., 2020). Some groups have taken unbiased approaches to comprehensively study motor function (Lipka et al., 2016) or used yeast 2 hybrid systems to identify RBP-kinesin interactions at higher throughput scale (Lang et al., 2021). Here, we have not only augmented the known set of RBP-kinesin interactions, but also described methods for how to identify and characterize additional pairs, including dissection of modular RBP domains and their distinct roles in subsequent steps of post-transcriptional regulation.

MBNL proteins bind mRNA targets to localize them to myoblast membrane fractions, immature neurites, and brain neuropil (Taliaferro et al., 2016; Tushev et al., 2018; Wang et al., 2012), but underlying mechanisms of this process have been unclear. Here, we show that MBNL interacts with specific motor proteins, Kif1bα and Kif1c, for directed transport (**Fig. 2**). No RBPs were previously known to specifically interact with these motors. Surprisingly, we found that these interactions are mediated through zinc finger domains in a non-RNA binding-dependent manner, with each pair of zinc fingers sufficient for kinesin association (**Fig. 4B-D**). This observation prompted us to assess potential interactions with proteins containing similarly structured ZnFs, such as Tis11d. Although Tis11d binds AU-rich motifs rather than YGCY, its ZnFs fold similarly to those of MBNL1 (Teplova and Patel, 2008). Indeed, Tis11d recapitulated similar patterns of kinesin specificity, showing recruitment by Kif1b⍺ and Kif1c, but not Kif1bβ (**Fig. 4E, F**). It remains to be tested whether other RBPs with similar ZnF structures show comparable profiles of kinesin specificity, including ZC3H14, which regulates polyA tail length and synaptic protein expression (Rha et al., 2017), and CPSF4, a core polyadenylation and cleavage factor that recognizes the AAUAAA polyadenylation signal in mRNA (Shimberg et al., 2016). Additional studies would be required to determine the precise nature of kinesin-MBNL association, and whether additional protein adapters are required to form, stabilize, or disrupt these interactions. Overall, our findings further raise the possibility of a broader, well-defined code that dictates RBP-kinesin specificity, with implications for understanding and manipulating microtubule-dependent RNA localization patterns.

Kif1bα has been implicated in mitochondrial trafficking (Nangaku et al., 1994; Wozniak et al., 2005) and localizes to postsynaptic densities (Mok et al., 2002), but was not previously known to localize RNAs or associate with RBPs. The alternatively spliced isoform Kif1bβ has been more extensively studied. Kif1bβ plays roles in transport of synaptic vesicle precursors through IGF1R, regulates axonal outgrowth, and is implicated in Charcot-Marie-Tooth disease and neuroblastoma (Xu et al., 2018; Zhao et al., 2001). Kif1c, whose tail domain is 58% similar to Kif1bα (compared with 46% similarity to Kif1bβ and 21% similarity to Kif5b), is involved in transport of mitochondria and dense core vesicles, and may be the fastest kinesin (Dorner et al., 1998; Lipka et al., 2016). Although it is plus-end-directed, it undergoes bidirectional transport through association with dynein and is subject to an autoinhibitory mechanism in which its stalk inhibits the motor domain in the absence of Hook3 and Ptpn21 (Kendrick et al., 2019; Siddiqui et al., 2019). Kif1c transports mRNAs, including its own transcript, to cell protrusions through an APC-dependent mechanism (Costa et al., 2020; Pichon et al., 2021). RNA binding proteins involved in Kif1c-mediated RNA transport were not known previously to our study. It remains to be determined whether MBNL and RNPs modulate motor activation or transport properties. Notably, at the transcript level, Kif1bα and Kif1c together comprise ∼75% and ∼50% of the total kinesin transcript pool in muscle and heart tissue, respectively, and ∼26% and ∼30% in the substantia nigra and spinal cord, respectively (**Table S3**). These high abundances might imply particularly important roles for these motors to transport cargoes in these tissues.

From these studies, it is unclear whether a single ZnF pair can simultaneously associate with binding targets. Differential RNA binding properties of each ZnF pair have been characterized, with ZnF(1,2) showing greater specificity than ZnF(3,4) (Hale et al., 2018). In our assays, we did not detect strong differences in kinesin association between ZnF(1,2) versus ZnF(3,4) (**Fig. 4D**). Thus, one possibility could be that one ZnF pair engages kinesin, while the other pair engages RNA. Alternatively, both pairs of ZnFs could engage RNA simultaneously with a single kinesin tail, or potentially even 2 separate kinesin molecules. Given the dimerization potential of MBNL1 via exon 7 (Tran et al., 2011), multivalency for both RNA binding and kinesin association can be achieved, with a total of 4 ZnF pairs linked together in a single complex. Indeed, we observed accumulation of Ncl mRNA, an MBNL target, at CUG RNA foci in both N2A and CAD cells (**Fig. 3F**, **Fig. S4**), raising the possibility that multiple RNAs could be tethered through 2 distinct MBNL ZnF pairs, or via dimerized MBNL proteins. Indeed, expression of MBNL lacking the C-terminal tail lessens the stability of CUG RNA foci (Arandel et al., 2022). Given that kinesin associates with MBNL ZnFs, the potential for tethering may not only apply to RNAs but also motors – since kinesins and dyneins work in teams to carry protein complexes and/or vesicles (Encalada et al., 2011; Hendricks et al., 2010), RBPs or RNPs could serve as multivalent adapters to bring multiple motors together into a single complex.

RNA localization has been proposed to involve not only RNP assembly and engagement with motors, but also anchoring to certain destinations. The sushi-belt model proposes that “hungry synapses” remove RNPs from the belt when activated (Doyle and Kiebler, 2011). It remains unclear how RNAs anchor at synapses, although interactions with intracellular membranes, including ER, is one possibility. MBNL targets are enriched in signal peptides, and many encode membrane proteins (Wang et al 2012). ER is enriched at dendritic branch points and synapses (Cui-Wang et al., 2012), which may also serve as sites of docking for MBNL-containing RNPs. Although we did not anticipate that a specific region of MBNL itself would be sufficient to engage membranes and anchor mRNAs, we found that the MBNL1 C-terminal domain alone is sufficient to slow RNP movement, limit total distance traveled, and tether RNAs to triton-sensitive fractions of mouse myoblasts. Unstructured, intrinsically disordered, and low complexity domains can enhance protein-protein or protein-RNA interactions and facilitate the formation of phase-separated condensates (Van Treeck and Parker, 2018). Interestingly, another RBP helps form an ER-associated, membraneless, “TIGER” domain, Tis11b (Ma and Mayr, 2018). Therefore, not only do MBNL and Tis11 family members share preferences for certain kinesins via their ZnFs, but may also exhibit parallels in membrane anchoring activities. It remains to be determined whether Tis11 and MBNL family members work together to recruit RNAs to ER, or alternatively, drive distinct cellular processes.

The association of RBPs with membranes also has implications for RNA localization by way of motile endosomes and lysosomes. Tethering of RNA granules to endosomes in a Rab7a-dependent manner (Cioni et al., 2019) or to lysosomes in an ANXA11-dependent manner (Liao et al., 2019) may hold implications for MBNL-mediated anchoring of RNAs to membranes. Similar to MBNL, multiple domains of ANXA11 perform distinct functions, with a low-complexity domain being essential for phase separation with membraneless RNA stress granules and the C-terminal annexin-repeat region acting as a binding partner for lysosomes. Mutations in the low complexity domain and annexin domains of ANXA11 have also been associated with ALS, highlighting the importance of unstructured domains and membrane tethering for cellular function and homeostasis (Nahm et al., 2020; Smith et al., 2017). It remains to be determined whether MBNL also tethers RNAs to endosomes or lysosomes, and whether there is specificity for certain types of vesicles as defined by Rab composition. MBNL is also post-translationally modified (Wang et al., 2018), however, it is unclear if modification is involved in MBNL-membrane associations, either through an adapter or a direct lipid insertion. Together, these observations illustrate the modularity of both globular and unstructured domains of RBPs to carry out all steps of post-transcriptional regulation, and the potential for multiple overlapping mechanisms to co-exist in the cell. Multiple neurological, neurodegenerative, and neuromuscular diseases are caused by mutations or perturbations to RBPs and kinesins. Mutations in highly expressed RBPs such as hnRNP A1, hnRNP A2B1, TDP-43, and FUS are associated with ALS (reviewed in Kapeli et al., 2017).

Mutations in the ATP binding site of KIF1Bβ are linked to Charcot-Marie-Tooth disease type 2A (Zhao et al., 2001), motor or tail domain of KIF1C cause autosomal recessive hereditary spastic paraplegia or ataxia (Caballero Oteyza et al., 2014; Dor et al., 2014), motor or stalk domains of KIF5A cause hereditary spastic paraplegia, and tail domain of KIF5A cause ALS (Ebbing et al., 2008; Nicolas et al., 2018), the last of which may be explained by disruption to auto-inhibition (Pant et al., 2022). Depletion of SMN in spinal muscular atrophy perturbs RNP granule formation and has deleterious effects on local translation in motor neurons (Fallini et al., 2011; Donlin-Asp et al., 2017). In DM1, MBNLs are depleted by expanded CTG repeat expression, with clear changes in global splicing patterns and potential changes in RNA localization in myoblasts (Wang et al., 2012). Although implied by studies of cells depleted of MBNLs by RNAi, our observations here confirm that expression of expanded CTG repeats impairs RNA localization in neural cells. Taken together, these findings prompt considerations about other diseases that could be exacerbated by compromised transport, including Huntington’s disease, Alzheimer’s disease, and other dementias. Here, we explored functional consequences of MBNL sequestration and kinesin perturbation by dominant negative tail expression in N2A and CAD neurites. As might be expected, we found modest overlap in shared targets of MBNL, Kif1bα, and Kif1c, given that these kinesins likely also associate with many other RBPs. Nucleolin mRNA was confirmed as a *bona fide* target that relies on these players; it encodes an RBP that regulates ribosome biogenesis but also localizes to distal neurites (Abdelmohsen et al., 2011; Perry et al., 2016). Ncl protein retrogradely transports importin β1 mRNA back to the soma to control cell size and associates with Kif5a through its glycine/arginine-rich domain (Doron-Mandel et al., 2021; Perry et al., 2016). Here, we find that Ncl mRNA itself is localized by MBNL and Kif1bα/Kif1c, raising the possibility that Ncl protein in neurites may arise via local translation. Indeed, MBNL1 CLIP tags appear throughout the Ncl mRNA, and it may be translationally repressed while transported by kinesins. Thus, this circuit may control cell size during neurogenesis and potentially even modulate susceptibility to cancer, as mutations connected to KIF1B have been implicated in neuroblastomas (Yang et al., 2001), paragangliomas, and pheochromocytomas (Evenepoel et al., 2017).

The observation that MBNL protein activity influences the abundance of Kif1bα raises the possibility of feedback between nuclear and cytoplasmic processes. It remains to be determined whether MBNL directly regulates the splicing or polyadenylation of Kif1b pre-mRNA, but an attractive possibility is that this architecture allows a single RBP to control and buffer the number of transport vehicles it can employ to carry out all of its functions. During muscle differentiation, MBNL activity rises, Kif1bα abundance increases, and there is overall increased reliance on microtubules to distribute mRNAs throughout the cytoplasm (Denes et al., 2021). Conversely, depletion of MBNLs in DM not only impairs localization functions of MBNLs, but also reduces the total pool of Kif1bα available for transport. MBNL CLIP tags also appear in the Kif1c 3’ UTR, whose mRNA is also localized to neurites and has multiple alternative polyadenylation sites (Batra et al., 2014; Costa et al., 2020). An open question is whether MBNL may also regulate Kif1c RNA processing and local translation, thus controlling expression of an additional transport partner. Finally, although KIF13A did not recruit MBNL in our kinesin tail screen, exon 32 is believed to be an MBNL splicing target and is a robust biomarker of disease status in DM1 – functions of exon 32 are unknown, and a potentially fruitful line of investigation may be to probe its roles in the pathways discussed here (Nakamori et al., 2013; Venables et al., 2013).

Finally, we establish methods for studying kinesin-RBP-mRNA interactions. Our implementation of the centrosome recruitment assay further augments a robust, well-established method, and could reveal many previously unknown kinesin-RBP interactions if applied to expanded targets. RNA-MRI reveals RBP properties by directly tethering RBP domains to the MS2 coat protein and assembling them onto an MS2 reporter RNA, painting a “dynamic caricature” of the RBP. Our assay has similarities to the “TnT translation and tether” assay (Cialek et al., 2022), but uses Halo signal to directly image RNPs that contain the RBP of interest. Inducible components yield signal to noise properties required to generate high quality movies amenable to automated computational analyses and could be expanded to many additional RBPs. The techniques outlined here provide a platform for further investigation of kinesins, RBP domains, and mRNAs that influence transport, with implications for our global understanding of mRNA localization, as well as how they may go awry in genetic diseases including DM.

## Contributions

R.P.H. designed plasmids and performed transfections, stable cell line creation, microscopy, and data analysis related to the centrosome recruitment assay, neurite fractionation, MEF fractionation, and MCP-Halo experiments. K.R.M. performed all experiments related to the split kinesin assay, immunofluorescence in primary neurons, and live neuronal imaging of MBNL granules. A.J.K. cloned full-length mScarlet kinesin plasmids and performed live imaging and data analysis of MBNL-kinesin co-transport in live neurons. L.K. performed co-immunoprecipitation experiments of MBNL1 with full-length kinesins and fixed colocalization experiments with endogenous kinesin and MBNL1. L.T.D. and T.S participated in plasmid construction and experimental design. D.P.B. and Z.L. performed experiments involving transfection and data analysis of centrosome recruitment with single MBNL1 ZnF mutants. K.L. assisted with study design. G.J.B and E.T.W. conceived of the project, supervised and assisted in the design of all study components, and provided resources and funding. R.P.H., G.J.B. and E.T.W. wrote the manuscript.

## Acknowledgements

We dedicate this manuscript posthumously to Kun Lin, graduate student at Emory University. We thank Gary Banker for providing kinesin tail and cargo-expressing plasmids used in the split kinesin assay and Andy Berglund (University at Albany) for providing MBNL1 plasmids with RIM and CM mutations. We thank Edouard Bertrand (Institut de Génétique Moléculaire de Montpellier) for the donation of a plasmid expressing a 45xMS2 array that was used to design the inducible RNA reporter. We also thank Maury Swanson (University of Florida) for donating MBNL1/2 double knockout fibroblasts and Chris Burge (Massachusetts Institute of Technology) for donating N2A and CAD cells. An additional thanks to Arielle Valdez-Sinon for technical assistance with co-immunoprecipitation experiments, Kendra McKee for technical assistance to prepare RNAseq libraries, Hailey Olafson for RNA-seq data processing, and Chase Kelley for assistance with particle tracking analysis. We thank Belinda Pinto, Chase Kelley, and other members of the Wang and Bassell labs for insightful comments and suggestions. Finally, we thank the Emory University Microscopy Core and University of Florida Center for Neurogenetics for use of microscopy equipment in fixed and live cell imaging experiments. L.K. was supported by NIH NRSA training grant F31 NS117086. This work was supported by an NIH R01 NS114253 awarded to G.J.B and E.T.W., and a Ben Barres Early Career Acceleration Grant from the Chan Zuckerberg Initiative awarded to E.T.W.

## Methods

### Plasmids and Cloning

All plasmid fusions assembled in-house were generated through standard ligation-based methods using the In-Fusion cloning kit according to manufacturer protocols (Takara, 638920). MBNL1 RNA-interaction and chelation mutant sequences were designed based on plasmids generated by Andrew Berglund (Purcell et al., 2012). Tis11d coding sequence was synthesized by IDT and Rbfox2 was amplified from mouse cDNA. All above sequences along with MBNL1 and MBNL2 isoforms imaged in primary neurons were cloned downstream of GFP in a pEGFP-C1 plasmid backbone under control of a CMV promoter (Takara). Full-length kinesin sequences were amplified from mouse cDNA and inserted upstream of an mScarlet-containing plasmid in a pcDNA3.1 backbone under the control of CMV promoter.

For dominant negative conditions, kinesin tail sequences were amplified from mouse cDNA and cloned into pEGFP-C1 as described above. For the centrosome recruitment assay, tail sequences were inserted downstream of a BicD2 dynein adapter in pcDNA3.1 under control of a CMV promoter. Plasmids containing FRB split kinesin tails and FKBP motors were a gift from Gary Banker and described previously (Bentley et al., 2015; Yang et al., 2016). Plasmids carrying exons 11-15 of the DMPK gene expressing either 0 or 480 CTG repeats under the control of a CMV promoter were a gift from Tom Cooper (Baylor College of Medicine).

PB-Neo-pERV3 contained a bicistronic ecdysone receptor expression cassette under control of EF1a promoter and PGK-driven neomycin resistance cassette was first stably introduced into C2C12 myoblasts. An MS2 reporter plasmid (PBPuroPonA-56xSunTag-45xMS2) containing a 56x SunTag coding sequence with a minimal intron upstream of 45x MS2 hairpins in the 3’ UTR and downstream of an ecdysone-responsive promoter with a blasticidin resistance cassette under control of a PGK promoter was then introduced. Finally, plasmids containing tandem MCP with nuclear localization sequences were cloned downstream of a tetracycline-responsive promoter with a puromycin resistance cassette under control of a PGK promoter. An MCP sequence was derived from a plasmid deposited by Robert Singer in the Addgene plasmid repository (Addgene, 40649). Subcloning into the MCP plasmid sequentially introduced the HaloTag sequence (Promega) and MBNL fusions (PB-PuroTet-MCP-HaloTag). MBNL1-GFP isoform sequences and mutants were amplified from previously established plasmids (see above) and were designed for stable expression of nuclear and cytoplasmic MBNL1 in C2C12 myoblasts and mouse embryonic fibroblasts (MEFs). MS2 array sequence is derived from a plasmid gifted by Edouard Bertrand. All transgenes for inducible MS2-MCP system and stable expression of MBNL1-GFP isoforms or mutants were flanked by PiggyBac transposon arms (Li et al., 2013; Wyborski et al., 2001).

### Cell Lines and Transfection

Neuro2a cells (N2A; gift from Chris Burge), MBNL double-knockout MEFs (DKO MEFs; gift from Maury Swanson), and C2C12 myoblasts (ATCC, CRL-1772) were cultured in DMEM supplemented with 100 U/mL penicillin, 100 mg/mL streptomycin, and 10% fetal bovine serum. CAD cells (gift from Chris Burge) were cultured in 1:1 DMEM:Ham’s F12 supplemented with 100 U/mL penicillin, 100 mg/mL streptomycin, and 8% fetal bovine serum. All cells were kept in a humidified incubator with 5% CO2 at 37°C. To induce N2A and CAD cell differentiation, media was changed to DMEM without serum.

For the split kinesin assay, N2A cells were transfected with Lipofectamine 2000 (Thermo Fisher, 11668027) according to manufacturer’s instructions, then imaged after overnight expression and ∼3 hour incubation with 500 nM A/C heterodimerizer linker drug (Takara, 635055) or 100% ethanol. 10μM S-Trityl-L-cysteine was added shortly after transfection for BicD2 experiments to stall cells in interphase and stabilize centrosomes. All other transfections in N2A, CAD, C2C12, and MEF lines were performed using TransIT X2 (Mirus, MIR6003) according to manufacturer’s instructions.

### Production of Stable C2C12 Myoblasts and MEFs

To make an MCP-MS2 reporter line, first, a wild-type C2C12 myoblast line was co-transfected with PB-Neo-pERV3 and an mPB plasmid expressing a PiggyBac transposase. PiggyBac allows for stable integration into the host cell genome (Li et al., 2013). C2C12 myoblasts were then selected with 500 µg/mL G418 (Geneticin) for 2 weeks. Following recovery, C2C12s were then transfected with PB-PuroPonA-56xST-45xMS2 for stable integration and selected with 10 µg/mL blasticidin for 1 week. Following a second recovery, C2C12 myoblasts were transfected with PB-PuroTet-MCP-HaloTag alone or fused to various MBNL1 proteins for stable integration and selected with 5 µg/mL puromycin for 1 week.

Other C2C12 myoblasts expressing various MBNL1 isoforms or DKO MEFs expressing MBNL1-41 and truncated mutants in PiggyBac plasmids were transfected similar to above and selected with 5 µg/mL puromycin for 1 week. All transfections for stable cell lines were performed using TransIT X2 (Mirus, MIR6003) according to the manufacturer’s instructions.

### Primary Mouse Cortical Neuron Culture, Transfection, and Transduction

Cortices isolated from E17 mouse embryos were dissected and dissociated with trypsin at 37°C (Thermo Fisher Scientific, 15050-065). Cells were plated in MEM/FBS on coverslips previously coated with 1mg/ml poly-L-lysine in Borate Buffer for 72 hours followed by three 1 hour washes with sterile water. After 2 hours, medium was replaced with astrocyte conditioned Neurobasal Plus Medium (Gibco, A3582901) supplemented with B-27 Plus Supplement (Gibco, A3582801) and GlutaMAX Supplement (Gibco, 35050061).

For live imaging of GFP-MBNL-kinesin co-transport in primary cortical neurons, cells were transfected with Lipofectamine 2000 (Thermo Fisher, 11668027) as follows: fresh culture medium was added to the cells 1:1, then half of the medium was collected. DNA/Lipofectamine mix was prepared according to manufacturer’s instructions and left with the cells for 1.5 hours, after which medium was replaced with the collected medium. Neurons were imaged after overnight expression following incubation with 100 ng/mL BDNF for 30 minutes. Live cell imaging of GFP-MBNL1 and GFP-MBNL2 isoforms involved transfection with Neuromag (OZ Biosciences, NM50200) at 7 DIV according to the manufacturer’s protocol. Cells were incubated on a magnet at 37°C for 20 minutes, incubated with 100 ng/mL BDNF after 18-24 hours, then imaged after 30 minutes. 8 DIV cortical neurons were transduced with adeno-associated virus containing ssDNA encoding HA-MBNL1. Four days later, at 12 DIV, the cells were processed for quantitative colocalization imaging of HA-MBNL1 and endogenous KIFs.

### Immunofluorescence (IF)

Cells were washed once with phosphate-buffered saline (PBS) then fixed with 4% paraformaldehyde (PFA) for 10 minutes at room temperature (RT). Cells were then washed 3X with PBS and permeabilized with 0.2% Triton X-100. Cells were washed again 3X with PBS (or Tris-Glycine buffer for neuronal staining) and incubated with primary antibody diluted in PBS containing 3% bovine serum albumin (5% for neuronal staining) and 0.1% Tween-20 (AB buffer) for 1 hour at RT. Primary antibodies used for IF include: mouse ɑ-MBNL1 (1:100, Developmental Studies Hybridoma Bank, MB1a(4A8)), mouse ɑ-MBNL2 (1:100, Developmental Studies Hybridoma Bank, MB2a(3B4)), rabbit ɑ-Map2 (1:1000, Millipore-Sigma, AB5622), guinea pig chicken ɑ-Map2 (1:1000, Synaptic Systems, 188 004), ɑ-Tau (1:1000, Aves Labs, TAU), mouse ɑ-Myc (1:100, Developmental Studies Hybridoma Bank, 9E10), mouse ɑ-γ-Tubulin (1:500, Abcam, ab11316), rabbit ɑ-FLAG (1:1000, Cell Signaling Technologies, D6W5B), mouse ɑ-KIF1B – recognizes KIF1Bβ (1:50, EMD Millipore, MABC309), mouse ɑ-KIF1B - recognizes KIF1Bα (1:50, Santa Cruz, sc-376246), rabbit ɑ-KIF1C (1:200, Abcam, ab125903), rabbit ɑ-HA (1:800, Cell Signaling Technologies, C29F4), and mouse ɑ-HA (1:1000, BioLegend, 901501). Cells were then washed 3X with PBS and incubated with secondary antibody in AB buffer (1:500, ɑ-mouse Alexa Fluor 488; 1:500, ɑ-mouse Alexa Fluor 647; 1:1000, ɑ-rabbit Alexa Fluor 647; 1:500, ɑ-chicken 647; 1:500, ɑ-guinea pig 647) for 1.5 hours at RT. Cells were washed 3X with PBS with DAPI (1 ng/uL) included on the final wash, mounted, and imaged.

### Western Blotting

For MEFs cytosolic and membrane lysates, protein concentrations were quantified using Pierce BCA protein assay kit (Thermo Fisher, 23225), processed by diluting in 4X LDS buffer (Thermo Fisher, NP0008), and separated on 4-12% Bis-Tris polyacrylamide gels (Thermo Fisher, NP0336) after denaturation for 5 minutes at 95°C. Gel electrophoresis was performed for ∼90 minutes at 100 V in 1X MOPS running buffer (50 mM 3-(N-morpholino)propanesulfonic acid (MOPS), 50 mM Tris base pH 8.0, 3.5 mM SDS, 1 mM ethylenediaminetetraacetic acid (EDTA)). Transfer was performed on activated polyvinylidene fluoride (PVDF) membranes (Bio-Rad, 1620264) using iBlot 2 dry transfer machine (Thermo Fisher). Membranes were blocked with SEA BLOCK (Thermo Fisher, 37527) for 30 minutes at RT. Membranes were then incubated with primary antibodies diluted in SEA BLOCK overnight at 4°C. Primary antibodies used for western blot include: mouse ɑ-Hsp90 (1:1000, Abcam, ab13492), rabbit ɑ-Calnexin (1:1000, Abcam, ab22595), chicken ɑ-GFP (1:5000, Abcam, ab13970), and rabbit ɑ-β-Actin (1:2000, Cell Signaling Technologies, 4967). The following day, primary solution was removed and membranes were washed 2X with PBS and 0.1% Tween-20 (PBST). Membranes were then incubated with secondary antibodies (1:5000, LICOR, IRDye 800CW donkey α-rabbit; IRDye 800CW donkey α-chicken; IRDye 680LT donkey α-mouse; IRDye 680LT donkey α-rabbit) diluted in SEA BLOCK for 1 hour at RT. Membranes were then washed again 2X with PBST and imaged using a LI-COR Odyssey CLx scanner at 700 and 800 nm excitation wavelengths.

### Immunoprecipitation

N2A cells were collected from 10 cm^2^ plates 48 hours after transfection of GFP-MBNL and kinesin plasmids and washed 2X with PBS. Cell pellets were resuspended with lysis buffer (10mM Tris-HCl pH 7.5, 150mM sodium chloride (NaCl), 0.5mM EDTA, 0.5% NP-40) containing protease/phosphatase inhibitor. Protein concentration was determined using Pierce BCA Assay (Thermo Fisher) and equal amounts of protein were incubated with 25 uL RFP-Trap magnetic agarose rotating for 1 hour at 4°C (Proteintech, rta-10). The beads were washed three times and boiled in SDS sample buffer for 10 minutes at 95°C. Proteins were separated on 4-20% Tris-Glycine polyacrylamide gradient gels (Bio-Rad, 4561096) and transferred onto 0.45mm nitrocellulose membranes. Membranes were blocked in 5% BSA diluted in PBS before being incubated overnight at 4°C with rabbit ɑ-MBNL1 antibody (1:1000, Millipore Sigma, ABE241) or mouse ɑ-RFP antibody (1:1000, Proteintech, 5F8). The following day, membranes were incubated with IRDye 680LT donkey α-mouse or IRDye 800CW donkey α-rabbit secondary for 1 hour at RT (1:20000, LICOR), along with Rhodamine-conjugated ɑ-GAPDH human Fab fragment (1:1000, Bio-Rad, 12004168). The blots were imaged on the ChemiDoc MP system and quantified using Image Lab software (Bio-Rad).

### Hybridized Chain Reaction Fluorescence *in situ* Hybridization (HCR FISH)

Probes, amplifiers, and buffers for performing HCR FISH 3.0 were purchased from Molecular Instruments. Probe sets constituted 10-30 probes, depending on target sequence. Cells were fixed with 4% PFA for 10 minutes at RT, washed 3X with PBS, then permeabilized with 0.2% Triton X-100. For combined IF/FISH experiments, cells were washed 3X with PBS, IF was performed first with 0.5% ultra-pure bovine serum albumin (Thermo Fisher, AM2616) and primary antibody in AB buffer for 1 hour. After washing 3X with PBS, secondary antibody incubation in AB buffer was performed for 1.5 hour followed by a second fixation in 4% PFA for 10 minutes at RT before proceeding with FISH. Cells were washed once with PBS then once with 2X saline-sodium citrate (SSC) and pre-hybridized with hybridization buffer (Molecular Instruments) for 30 minutes in a 37°C humidified chamber. Primary probes were diluted in hybridization buffer at a concentration of 1.2 pmol and incubated with cells for 16 hours in a 37°C humidified chamber. If repeat FISH was performed, a CAG10 Alexa Fluor 594 probe (Stellaris) was included in hybridization buffer at a concentration of 380 ng/uL. After primary incubation, cells were washed with probe wash buffer (Molecular Instruments) 4X in a 37°C humidified chamber, then twice with 5X SSC and 0.1% Tween-20 (SSCT), and finally pre-amplification was performed for 30 minutes at RT with amplification buffer (Molecular Instruments). During pre-amplification, appropriate HCR amplifier sets corresponding to primary probe initiators were heated separately in PCR tubes for 90 seconds at 95°C and allowed to cool to RT in the dark. Primed amplifiers were then added to amplification buffer at a concentration of 2.4 pmol and incubated with cells for 4 hours at RT. Cells were then washed 5X with 5X SSCT with DAPI (1 ng/uL) included on the final wash, mounted, and imaged.

### Fixed Cell Imaging

Results of HCR FISH in C2C12 myoblasts and N2A cells were imaged on a Zeiss LSM 880 Axio Observer microscope with Airyscan using a Plan-Apochromat 1.46 NA x100 oil objective and processed using ZEN software. Fixed cell imaging for centrosome recruitment assay was done with widefield illumination using a Plan-Apochromat 1.3 NA ×40 oil objective. Fixed cell imaging of the split kinesin assay in N2As and MBNL in primary neurons was performed using widefield illumination on a Nikon Eclipse TE300 inverted microscope with a Plan-Neofluar 1.4 NA x60 oil objective. For quantitative colocalization experiments, 12 images were taken in a z-series at 2um steps and deconvolved using a 3-D blind constrained iterative algorithm (AutoQuant, CyberMetrics). Imaris imaging software (Bitplane), namely the ‘Coloc’ module, was then used for analysis on the deconvolved images. Quantitative colocalization analysis involved creating a 3-D mask of the MAP2 channel from select neurites and excluding background signal from outside this volume. Within the MAP2 masked volume, two measures of colocalization, Pearson’s Correlation Coefficient and Mander’s Overlap Coefficient, were calculated between HA-MBNL1 and the respective endogenous KIF signals.

### Live Cell Imaging

All imaging in C2C12 myoblasts was performed on a Zeiss LSM 880 Axio Observer microscope with Airyscan using a Plan-Apochromat 1.46 NA ×100 oil objective. To obtain sufficiently high signal to noise ratios and resolve individual RNPs, we imaged cells in the low levels of ponasterone A and no doxycycline, relying on low and leaky expression from inducible promoters for MS2 and MCP components. Myoblasts were cultured on chambered coverglass (Thermo Fisher, 155409) with 2 uM Ponasterone A overnight and incubated with JF646 HaloTag ligand (Promega, GA1120) 15 minutes before imaging. Fluorobrite media (Thermo Fisher, A1896701) containing no phenol red supplemented with 100 U/mL penicillin, 100 mg/mL streptomycin, and 10% fetal bovine serum was used to wash and culture unbound HaloTag ligand and minimize autofluorescence. Cells were then placed in a humidified chamber with 5% CO2 at 37°C. Images were collected at 10 frames per second for 60 seconds. For imaging purposes, MCP-Halo expression was not induced by doxycycline due to noticeable, low-level expression observed in stable cells.

For live cell imaging of primary neurons, coverslips were transferred to a pre-warmed Chamlide CMB magnetic chamber (Quorum Technologies, CM-B18-1) and placed in warmed Hibernate E Low Fluorescence (BrainBits, HELF) CO2 independent media, supplemented with B27 Plus and GlutaMAX. Neurons were then supplemented with 100 ng/mL BDNF, a well-established paradigm used to induce transport of RNPs (Fallini et al., 2011; Zhang et al., 1999), and imaged after at least 10 minutes with using widefield illumination on a Nikon Eclipse TE300 inverted microscope with a Plan-Neofluar 1.4 NA x60 oil objective supplied with live cell imaging heating chamber. For imaging of motile GFP-MBNL isoforms, a 200 ms exposure time was used with images acquired every 260 ms for 1-2 minutes. Granules were manually tracked in ImageJ and velocities were calculated by dividing total time by total distance of each track. For imaging of co-transport with GFP-MBNL and kinesins, regions in either dendrites or axons were chosen where long processive movements (≥2 µm) were observed, then regions were converted to kymographs with ImageJ Reslice tool. Tracks were then labeled with a line ROI tool and measured. Distance and velocity were calculated on the basis of the line length and angle. Tracks that overlapped in both channels were categorized as co-transported particles.

### Centrosome Recruitment Analysis

N2A cells were transfected as described above with BicD2-KIF tail fusions and MBNL-GFP proteins at a ratio of 2:3 in 4-well chamber slides (Thermo Fisher, 154526). 24 hours after transfection, cells were fixed and processed for immunofluorescence as described above with a rabbit ɑ-FLAG antibody (1:1000, Cell Signaling Technologies, D6W5B) to visualize BicD2-KIF fusions. A collection of tile scans was taken from each condition on a Zeiss LSM 880 Axio Observer microscope with widefield illumination using a Plan-Apochromat 1.3 NA ×40 oil objective. After opening each tile scan image on ImageJ, cells and centrosomes were manually chosen, traced, and cataloged with the ROI Manager using MBNL-GFP and centrosomal FLAG signal as a mask, respectively. Care was taken to analyze cells that were not overexpressing tail proteins and did not have abnormally asymmetric centrosomes. Centrosome recruitment was quantified by taking the average intensity of MBNL signal at the centrosome divided by the average MBNL intensity in the whole cell.

### RNA Granule Dynamics

Time lapse images were captured using a Plan-Apochromat 1.46 NA ×100 oil objective at 10 frames/second. Background fluorescence was corrected and spot signal enhanced through setting a rolling ball radius of 3 pixels. Particle tracks were analyzed with the Fiji TrackMate software (Tinevez et al., 2016). RNP spots were identified based on approximate diameter (∼0.5 µm), consistent with previous observations of RNP granule size (Batish et al., 2012; Elvira et al., 2006), and quality across each time series. Spot track length threshold was set at >20 frames (2 sec). Spot location data was analyzed using custom scripts written in Python 3. Mean squared displacement (MSD) of each particle along the track was calculated at increasing time lags (up to lag time 7) and was used to quantify diffusion coefficients, which represents the slope of MSD plotted over lag time. Distance refers to the furthest point away from the track start a spot appears along the track.

### Neurite Fractionation and RNA-seq

Neurite fractionation was performed similar to a previously established protocol (Arora et al., 2021). The underside of hanging cell culture inserts with 1 µm pores (Corning, 353102) were coated with diluted Matrigel preparation (Corning, 354277) and placed into specialized, deep 6-well plates (Corning, 353502). 2 mL of 10% DMEM media was placed on top and underneath the hanging insert membrane prior to cell plating. N2A cells were then densely plated onto membranes at a concentration of 1×10^6^. After 24 hours, serum-containing media was replaced with no-serum media and neurites were allowed to grow for 48 hours. Subsequently, no serum media was removed from both sides of the membrane, which was washed once with PBS, then cell bodies were collected with a cell scraper in 1 mL PBS. Membranes were separated from their housing and placed into a lysis buffer (Zymo, R1060). Cell body and neurite samples from all wells of a 6-well plate were combined separately for each condition. After collecting all samples, cell bodies were spun down at 2,000 × *g* for 5 minutes and resuspended in 600 uL of PBS, with 100 uL reserved for RNA isolation. Reserved cell body volume was lysed, then cell body and neurite samples were processed for RNA isolation with a Quick-RNA Microprep Kit (Zymo, R1050). Following isolation, ribosomal RNA was depleted from 300 ng of the total RNA samples using the NEBNext rRNA Depletion Kit (NEB, E6310L), then processed for RNA-seq library prep with the NEBNext Ultra II Directional RNA Library Prep Kit (NEB, E7530L), and finally sequenced on an Illumina NextSeq 500. Fastq files were obtained using bcl2fastq2 v2.20. Gene expression values were quantitated by kallisto 0.43.0 (mapping to mm10 Refseq annotations) and normalized using the R “cyclicLoess” function. Localization ratios were computed by log2(neurite/soma). RNA-Seq data have been submitted to GEO (GSE207597).

### Membrane Fractionation

Extraction of cellular fractions was performed using lysis buffer (10 mM piperazine-N,N′-bis(2-ethanesulfonic acid) (PIPES), 0.25 M sucrose, 1 mM triethylene glycol diamine tetraacetic acid supplemented with 0.015% digitonin (cytosolic fraction) or 1% Triton X-100 (membrane fraction). Fractions were collected by incubating cells in 6-well plates with 500 uL cytosolic lysis buffer for 5 minutes at RT, collecting cytosolic sample, then incubating with membrane lysis buffer for 5 minutes at RT followed by collection of membrane fraction. Protein samples were processed by diluting in 4X LDS sample buffer (mentioned previously). RNA was isolated from the same samples using Direct-Zol RNA Miniprep Kit (Zymo, R2050) and iScript cDNA synthesis kit used to make cDNA (Bio-Rad, 1708890). Real-time amplification of the SunTag coding sequence, Gapdh (cytosolic), and Calnexin (membrane) using qPCR primers (IDT) with Q5 polymerase (New England Biolabs, M0492) was measured with a highly sensitive dsDNA detection dye (Lumiprobe, 11010) on a 96-well RT-PCR thermal cycler (Bio-Rad).

## Figures

**Figure S1.**
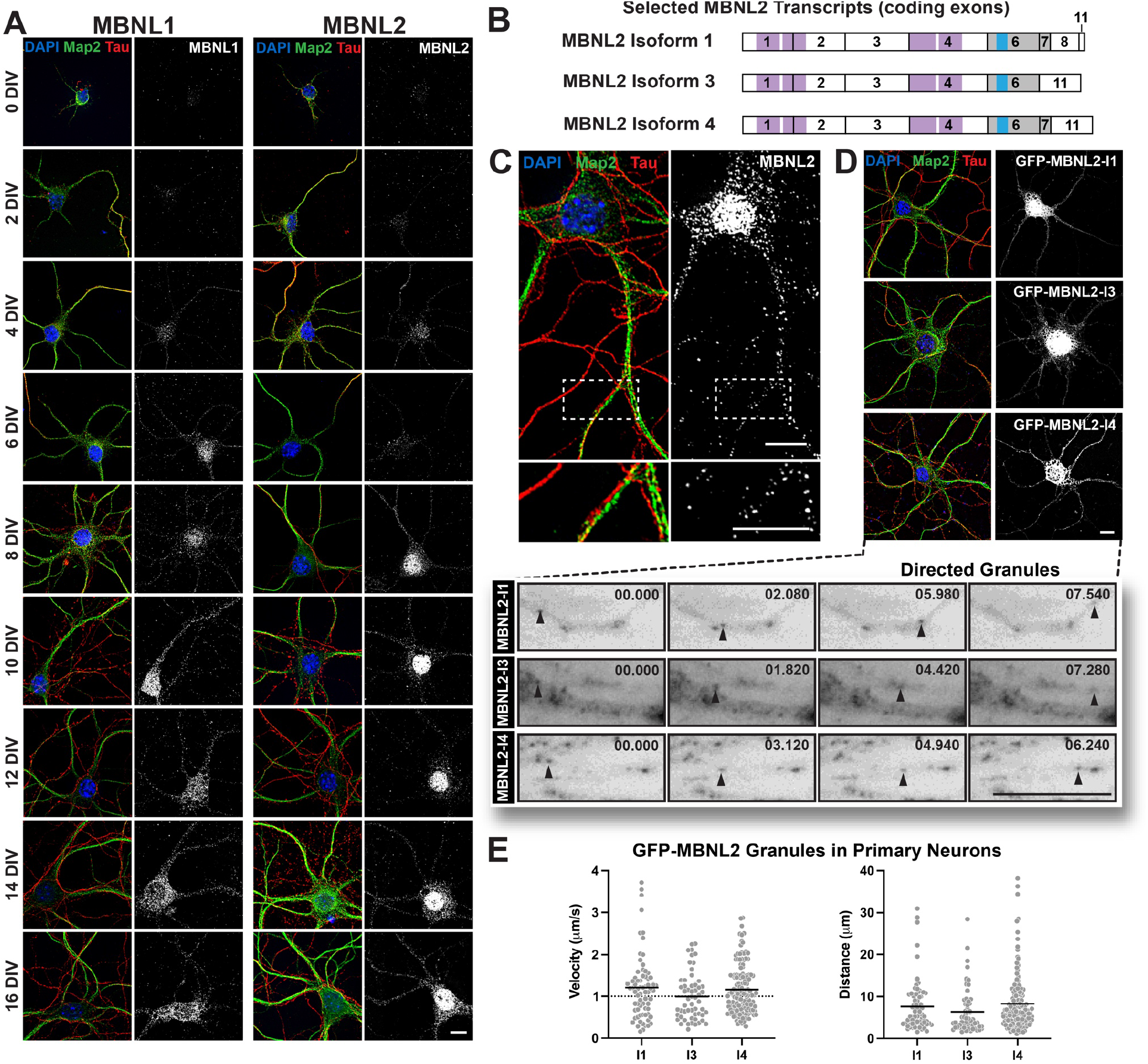
MBNL1 and -2 are developmentally regulated in primary cortical neurons and cytoplasmic MBNL2 isoforms are actively transported in axons and dendrites. A) Mouse primary cortical neurons were cultured for 0-16 DIV and fixed every other day. Immunofluorescence of MBNL1 and MBNL2 granules (white) in axons (Tau, red) and dendrites (Map2, green) corresponding to DIV. B) Exon and protein domain structure of cytoplasmic MBNL2 isoforms studied. nuclei (blue) labeled with DAPI. Scale bar = 10 µm. C) Representative image showing MBNL2 granules (white) in axons (Tau, red), dendrites (Map2, green), and nuclei (blue) of 9 DIV cultured primary mouse cortical neurons. D) Representative images of GFP-MBNL2-I1, -I2, and -I3 isoforms (white) in 9 DIV cultured primary mouse cortical neurons. Axons and dendrites labeled with Tau (red) and Map2 (green), respectively. Below are representative time lapse images of motile GFP-MBNL2 isoform granules in live mouse cortical neurons. Time (sec) indicated in top right. E) Quantitation of speeds and distances traveled by cytoplasmic MBNL2 granules (MBNL2-I1 – 70 tracks, 2.8 tracks/cell; MBNL2-I3 – 56 tracks, 1.87 tracks/cell; MBNL2-I4 – 141 tracks, 3.62 tracks/cell). Dotted line in velocity chart represents typical speed of microtubule-dependent transport. Bars represent mean. Nuclei labeled with DAPI (blue). Scale bars = 10 µm.

**Figure S2.**
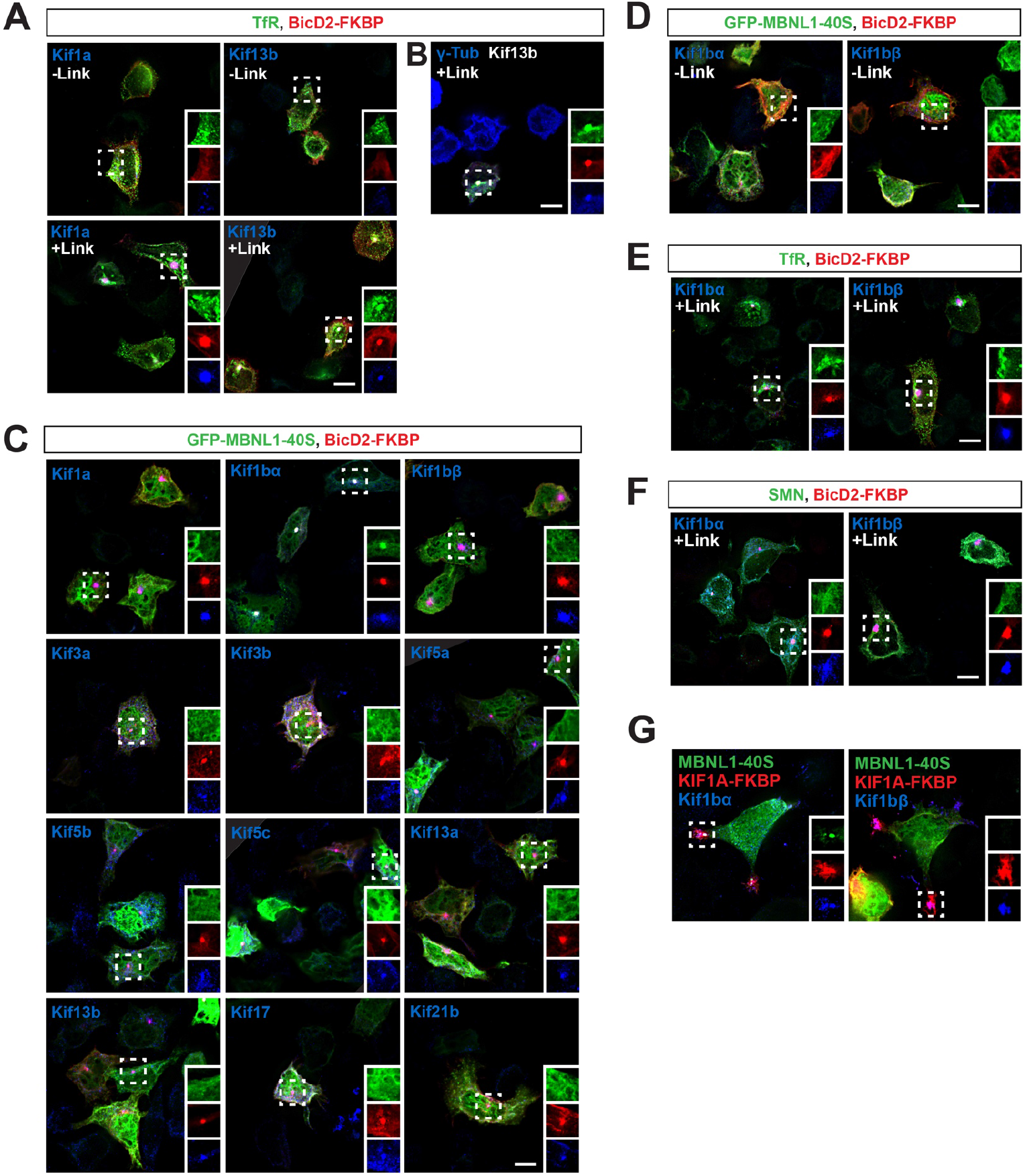
Kinesin tail screen with MBNL1 and controls. A) N2A cells were transfected with GFP-TfR (green), tdTomato-tagged BicD2-FKBP (red) and Myc-Kif1a-FRB or Myc-Kif13b-FRB in the presence of linker drug or 100% ethanol after ∼16 hours. Cells were fixed after 3 hours and processed for immunofluorescence with an A) Myc or B) γ-Tubulin antibody (blue). (C) N2A cells were transfected with GFP-tagged MBNL1-40S (green), tdTomato-tagged BicD2-FKBP (red) and Myc-tagged kinesin tails (Kif1a-FRB, Kif1bα-FRB, Kif1bβ-FRB, Kif3a-FRB, Kif3b-FRB, Kif5a-FRB, Kif5b-FRB, Kif5c-FRB, Kif13a-FRB, Kif13b-FRB, Kif17-FRB or Kif21b-FRB in the presence of linker drug. Cells were fixed after 3 hours and processed for immunofluorescence with a Myc antibody (blue). D) N2A cells were transfected with GFP-tagged MBNL1-40S (green), tdTomato-tagged BicD2-FKBP (red) and Myc-Kif1bα-FRB or Myc-Kif1bβ-FRB in the presence of 100% ethanol or E) linker drug. F) GFP-tagged SMN (green) was transfected with tdTomato-tagged BicD2-FKBP (red) and Myc-Kif1bα-FRB or Myc-Kif1bβ-FRB in the presence of linker drug. G) N2A cells were transfected with GFP-tagged MBNL1-40S (green), tdTomato-tagged KIF1A-FKBP (red) and Myc-Kif1bα-FRB or Myc-Kif1bβ-FRB in the presence of linker drug. Scale bars = 10 µm.

**Figure S3.**
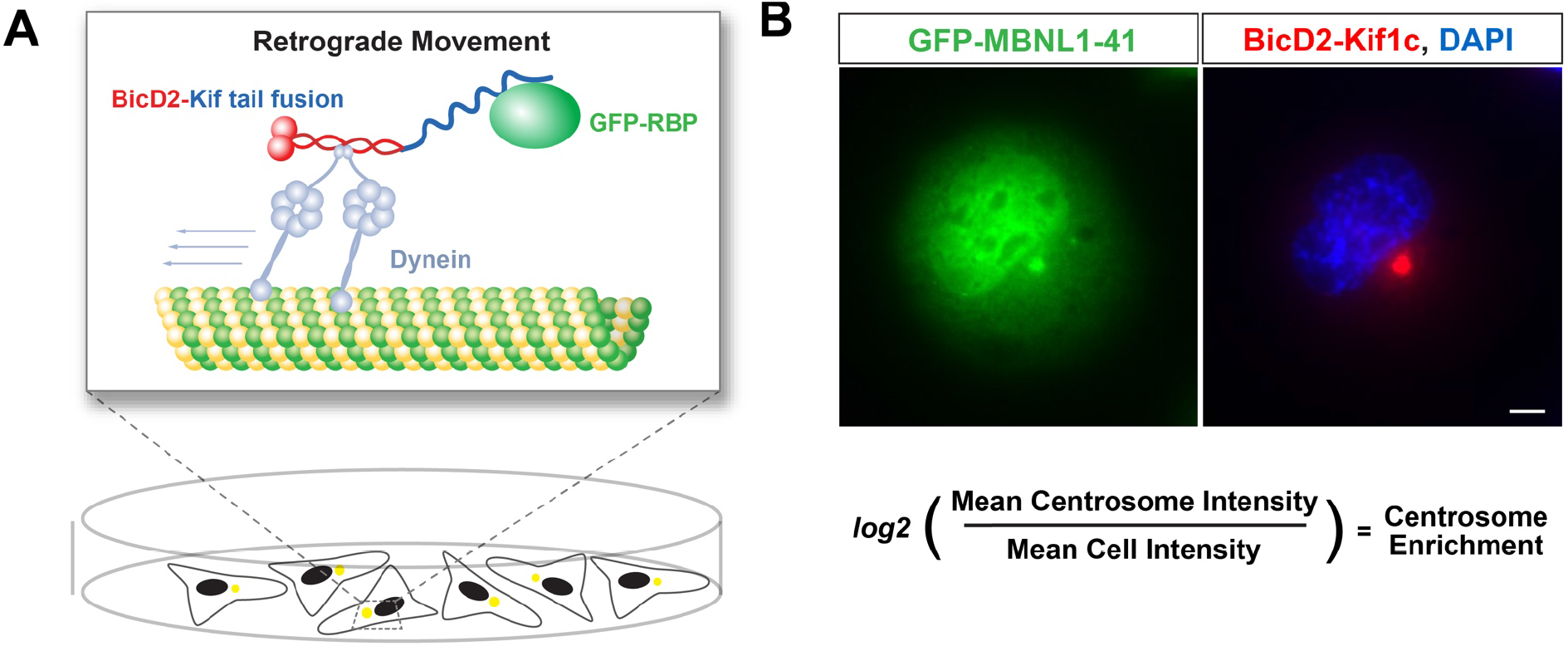
Schematic of the centrosome recruitment assay and quantitation. A) Schematic of the assay components and dynein engagement towards the centrosome. B) Representative images of a GFP-MBNL1-41 (green) when co-expressed with a FLAG-BicD2-Kif1c tail fusion (red). Nuclei (blue) labeled with DAPI. Scale bar = 5 µm. Centrosome enrichment is defined as the log2 value of the mean centrosome intensity divided by the mean cell intensity of the transported cargo.

**Figure S4.**
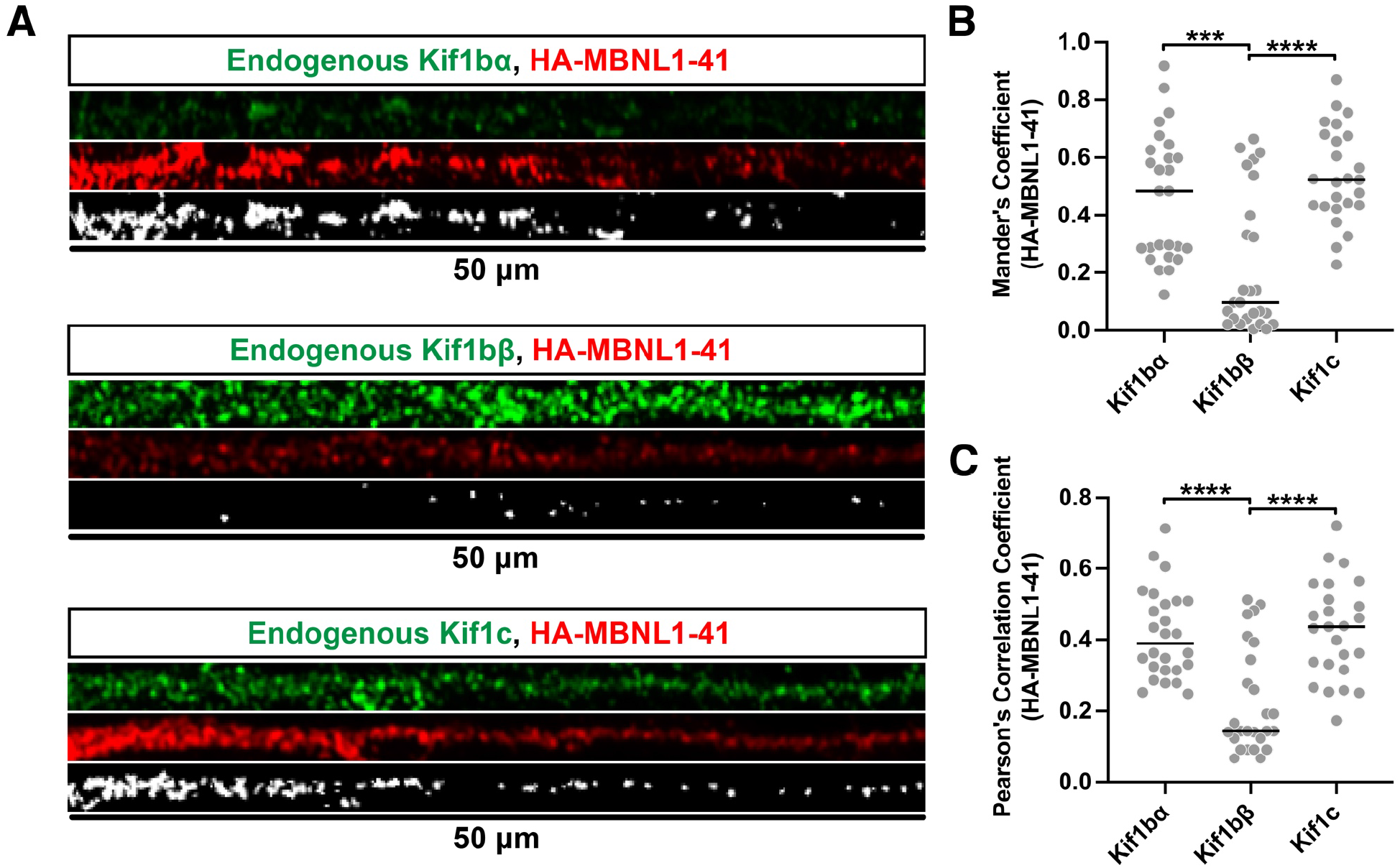
Endogenous Kif1bα and Kif1c co-localize with HA-MBNL1-41. A) Representative 50 μm neurite scans with immunofluorescence for endogenous kinesins (green) and transduced HA-MBNL1-41 (red) in 12 DIV primary mouse cortical neurons. Colocalized signal is represented below (white). B) Mander’s overlap coefficient of endogenous kinesins with HA-MBNL1. C) Pearson’s correlation coefficient of endogenous kinesins with HA-MBNL1-41. Kif1Bα = 26 neurites, Kif1bβ = 26 neurites, Kif1c = 25 neurites. Bars denote median. ***p<0.001, ****p<0.0001, One-way ANOVA with Tukey’s post-hoc test.

**Figure S5.**
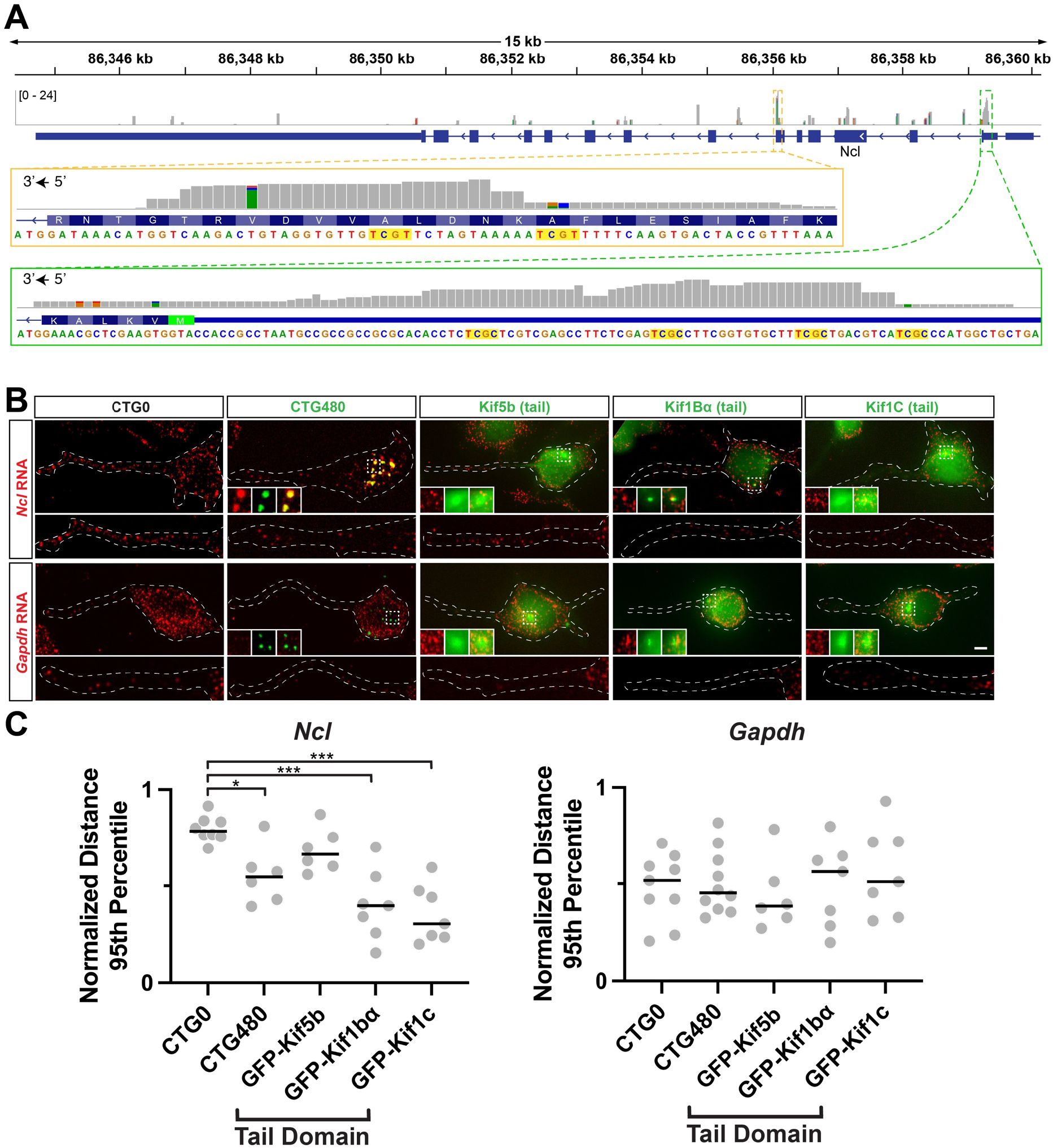
HCR FISH in differentiated CAD cells confirms Ncl mislocalization. A) MBNL1 CLIP from C2C12 myoblasts at the Ncl locus (mm10). Insets show two specific locations within the coding sequence (orange box) and 5ʹ UTR (green box), in which YGCY motifs are highlighted in yellow. B) Representative HCR FISH images for Ncl and Gapdh (red) with CTG repeats or dominant negative kinesin tails (green). Scale bar = 10 µm. C) Distances of Ncl and Gapdh HCR FISH spots from the soma of each cell across conditions. Distances were normalized as a fraction of total neurite length, and the 95th percentile of distances within each cell is plotted as an individual gray circle. Each cell (Ncl - CTG0: 10 cells, CTG480: 10 cells, Kif5b: 7 cells, Kif1bα: 7 cells, Kif1c: 7 cells; Gapdh - CTG0: 10 cells, CTG480: 10 cells, Kif15b: 6 cells, Kif1bα: 7 cells, Kif1c: 7 cells) contained >200 Ncl spots and >150 Gapdh spots. Bars represent the median across cells. *p<0.05, **p<0.01, Two-tailed Mann-Whitney U test.

