## Supplemental Information for "Muscleblind-like proteins use modular domains to localize RNAs by riding kinesins and docking to membranes"

### Supplementary Information

**Video S1. GFP-MBNL1 granules in live cultured neurons, related to Figure 1.** Cytoplasmic MBNL1-40 and -41 kDa isoforms were transfected by magnetofection into live cortical mouse neurons. Granules (black) exhibit directed motion. Particle track overlays seen in blue. 24 frames per second. Scale bars = 5 µm.

**Video S2. GFP-MBNL1 granules in live mouse embryonic fibroblasts, related to Figure 1.** Cytoplasmic MBNL1-40 kDa and -41 kDa isoform granules (white) stably expressed in live mouse C2C12 myoblasts. Granules exhibit mostly anchored behavior, with occasional diffusive or directed motion. 60 frames per second. Scale bars = 5 µm.

**Video S3. Representative MCP-Halo particle track output from TrackMate.** Cropped particle track output in TrackMate Fiji plugin from MCP-Halo RNP stably expressed in C2C12 myoblasts. 60 frames per second. Scale bar = 1 µm.

**Video S4. Representative particle tracks of MCP-Halo granules.** Representative movies of C2C12 myoblasts expressing MCP-Halo-MBNL1-RIM, -ΔC RIM, and -Δ3, ΔC RIM (left) with track overlays from TrackMate Fiji plugin (right). Particle tracks from full-length RIM exhibit a greater confinement and anchoring compared to ΔC RIM and Δ3, ΔC RIM that lack the MBNL1 C-terminal tail domain. 60 frames per second. Scale bars = 5 µm.

**Video S5. MCP-Halo protein and MS2 reporter are necessary to form visible granules.** Representative movies of C2C12 myoblasts expressing MCP-Halo alone or with the C-terminal tail domain (+C-terminus) and with (+MS2) or without (no MS2) a 45xMS2 reporter. Presence of the reporter is required to form visible RNP granules. 60 frames per second. Scale bars = 5 µm.

**Table S1.** TPM values from sequencing of neurite fractionation samples.

**Table S2.** Filtered $\log_2$ LR values of all conditions along with number of annotated MBNL1 CLIP sites in mouse brain.

**Table S3.** Kinesin transcript expression across tissues (average values for all samples in each tissue) as assessed by RNAseq from GTEX.
